# Using Deep Learning to Annotate the Protein Universe

**DOI:** 10.1101/626507

**Authors:** Maxwell L. Bileschi, David Belanger, Drew Bryant, Theo Sanderson, Brandon Carter, D. Sculley, Mark A. DePristo, Lucy J. Colwell

## Abstract

Understanding the relationship between amino acid sequence and protein function is a long-standing problem in molecular biology with far-reaching scientific implications. Despite six decades of progress, state-of-the-art techniques cannot annotate 1/3 of microbial protein sequences, hampering our ability to exploit sequences collected from diverse organisms. In this paper, we explore an alternative methodology based on deep learning that learns the relationship between unaligned amino acid sequences and their functional annotations across all 17929 families of the Pfam database. Using the Pfam seed sequences we establish rigorous benchmark assessments that use both random and clustered data splits to control for potentially confounding sequence similarities between train and test sequences. Using Pfam full, we report convolutional networks that are significantly more accurate and computationally efficient than BLASTp, while learning sequence features such as structural disorder and transmembrane helices. Our model co-locates sequences from unseen families in embedding space, allowing sequences from novel families to be accurately annotated. These results suggest deep learning models will be a core component of future protein function prediction tools.

---

Predicting the function of a protein from its raw amino acid sequence is a critical step for understanding the relationship between genotype and phenotype. As the cost of DNA sequencing drops and metagenomic sequencing projects flourish, fast and efficient tools that annotate open reading frames with function will play a central role in exploiting this data [1, 2]. Doing so will help identify proteins that catalyze novel reactions, design new proteins that bind specific microbial targets, or build molecules that accelerate advances in biotechnology. Current practice for functional prediction of a novel protein sequence involves alignment across a large database of annotated sequences using algorithms such as BLASTp [3], or profile hidden Markov models built from aligned sequence families such as those provided by Pfam [4, 5].

While these approaches are generally successful, at least one-third of microbial proteins cannot be annotated through alignment to characterized sequences [6, 7]. Moreover, the computational costs of methods such as BLASTp scale roughly linearly with the size of the labelled database, which is growing exponentially [8]. Broad protein families require multiple HMM profiles to model their diversity [9], while more than 22% of the highly-curated families in Pfam 32.0 have no functional annotation. More generally, models that predict function from sequence are limited by pipelines that require substitution matrices, sequence alignment, and hand-tuned scoring functions.

Deep learning provides an opportunity to bypass these bottlenecks and directly predict protein functional annotations from sequence data. In this framework, a single model learns the distribution of multiple classes simultaneously, and can be rapidly evaluated. Besides providing highly accurate annotations, the model’s intermediate layers can capture high-level structure of the data through learned representations [10] that can be leveraged for exploratory data analysis or supervised learning on new tasks with limited training data.

A number of recent papers have applied deep learning to achieve accurate protein function annotation using classification schemes such as GO terms and EC numbers [11–20], with some experimental validation [21], and also for DNA-protein interactions [22, 23]. The resulting learned data representations, also known as embeddings, for protein sequences also provide new exploratory tools with the potential for significant impact [19, 24–27]. However, existing deep learning work on protein function annotation often focuses on subsets of the protein universe, relies on additional information beyond primary sequence, or does not compare performance with existing state-of-the-art methods. These limitations reduce consideration of these approaches by the community.

In this paper, we ask whether deep learning models can complement existing approaches and provide protein function prediction tools with broad coverage of the protein universe, enabling more distant sequences to be annotated. We compare deep learning and existing approaches on the task of annotating unaligned protein domain sequences from Pfam-seed, which includes 17929 families, many of which have very few sequences [28]. For protein sequences, similarities between the test and train data mean it is essential to stratify model performance as a function of similarity between each held-out test sequence and the nearest sequence in the train set. We analyze both random and clustered splits, in which sequences are assigned to the test or train split based on cluster membership [29].

We find that the deep models make fewer errors at annotating held-out test sequences than state of the art HMM and BLASTp approaches across both the random and clustered splits. To confirm that the model has captured the structure of unaligned protein sequences, we use the joint representation learned across protein families in one-shot learning to annotate sequences from small families that the model was not trained on. These findings demonstrate that deep learning models can annotate proteins with their functions, and accelerate our ability to understand and exploit metagenomic sequence data.

## Results

In this section, we use the Pfam-seed dataset to construct benchmark annotation tasks, and compare the performance of deep learning models with existing alignment-based methods including BLASTp [30] and profile HMM methods [29]. Our stratified analysis of each data split takes into account the similarity between each held-out test sequence and the training set, and finds that the deep learning models perform well across the full range of sequence similarities. The benchmark includes a random train-test split^1^ of the 17929 Pfam families, where 80% of sequences are used for training (alignment retained for the profile HMMs only), 10% for model tuning (a ‘dev’ set) and 10% are held out as test sequences (Supplementary Table 1). Because protein sequences are related through evolution, there is a range of similarities between sequences in the train and test sets that confound the average reported accuracy. To address this, we stratify our analysis by the maximum percent identity of each test sequence with sequences in the train set.

Our CNN model (ProtCNN) significantly outperforms the alignment-based methods for sequences with between 30 and 90% maximum identity with the training set (p < 0.05, 2-sided McNemar test, Fig. 1A). The ProtENN model takes a simple majority vote across an ensemble of 13 ProtCNN models (Supplementary Fig. 1) to achieve an error rate of 0.16%, reducing both the HMM and BLASTp error rates by a factor of 9, to 201 misclassified sequences (Table 1). ProtENN is significantly more accurate than alignment-based methods for all sequence bins with less than 90% identity to the training set (p < 0.05, McNemar test, Fig. 1A). We implement profile HMM-based methods both by building a profile per family with `hmmbuild`, and then searching with `hmmsearch`, and also by using `phmmer` on unaligned sequences.

**Figure 1:**
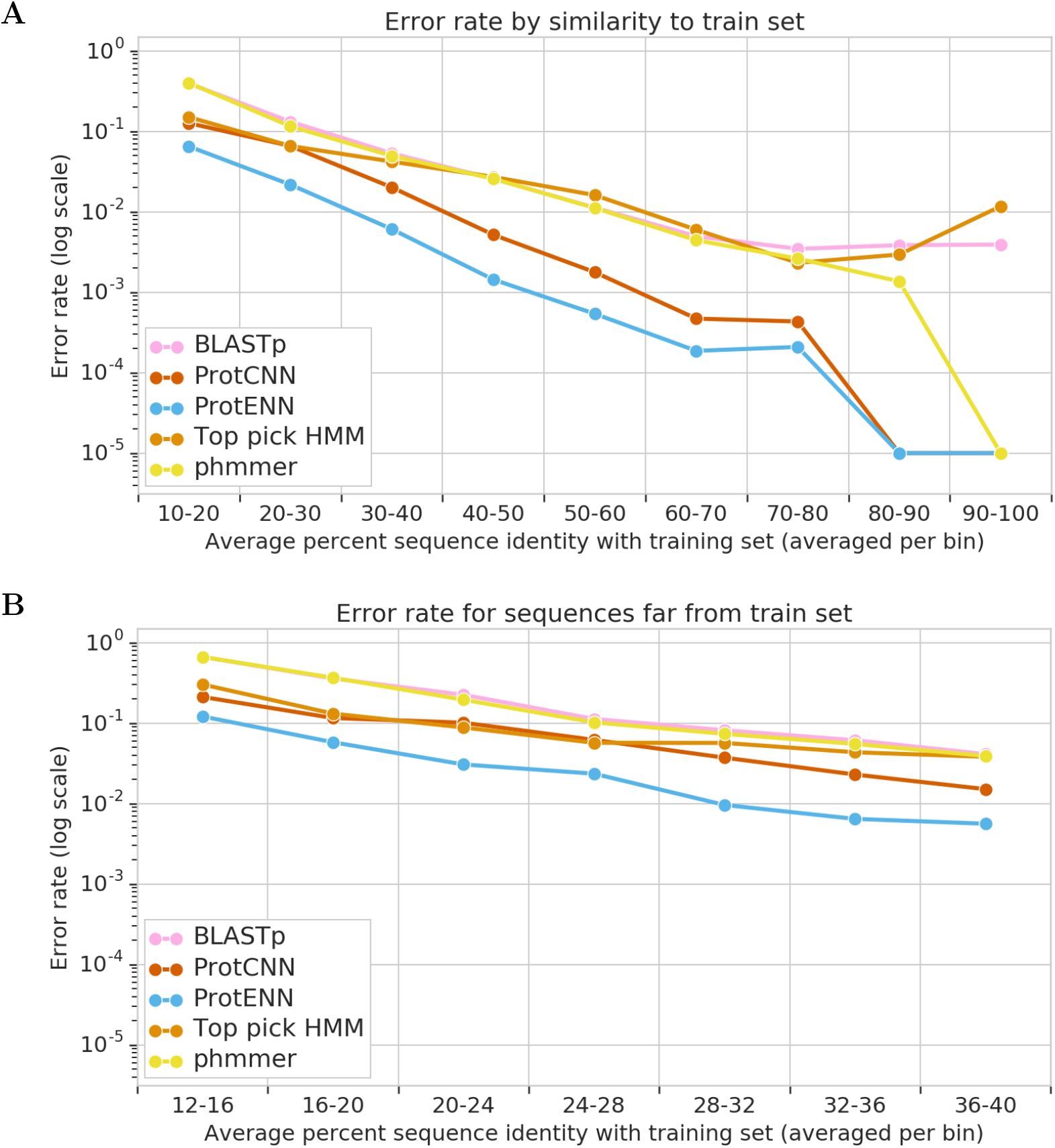
Model performance on the random split of Pfam-seed. (A) Held-out test error rate as a function of the maximum percent sequence identity across all training sequences. Top Pick HMM (Methods) is based on hmmsearch. Data is binned by maximum percent sequence identity with the training set; the x-labels describe the bin ranges. For each test sequence we use the Pfam-seed family alignment to calculate the percent sequence identity with the most similar train sequence in the same Pfam family (see Methods). ProtCNN makes significantly fewer errors than alignment-based methods for sequence identities in the range 30-90% and ProtENN is significantly better for sequence identities less than 90% (p < 0.05, McNemar test). (B) Zoomed plot of model performance for sequence identities below 40% (13457 sequences). Note that ProtENN is significantly better for all bins, including the 12-16% bin. The number of sequences per bin in either chart is available in Supplementary Tables 3 and 4.

**Table 1:** Performance on randomly-split data.

| Model | Error rate | Number of errors |
| --- | --- | --- |
| Top Pick HMM | 1.414% | 1784 |
| phmmer | 1.531% | 1932 |
| BLASTp | 1.654% | 2087 |
| k-mer | 9.994% | 12610 |
| RNN | 1.800% | 2271 |
| ProtCNN | 0.495% | 625 |
| **ProtENN** | **0.159%** | **201** |

The 13457 sequences most distant from the training set are further analyzed in Fig. 1B. We find that a single ProtCNN model displays similar performance to alignment-based methods across these remote sequences, while ProtENN obtains significantly greater accuracy across all bins. These results support the notion that ProtENN generalizes well to unseen parts of the data space available in the test split. We find 11 sequences that are consistently predicted incorrectly in exactly the same way by all ensemble elements of ProtENN. Supplementary Table 2 suggests that there is some ambiguity about their correct family labels (see also [31]). For example, our models predict that the sequence R7EGP4_9BACE/13-173 from Pfam family DUF1282, actually belongs to the YIP1 family. The `hhsearch` [4] tool predicts that DUF1282 is similar to the YIP1 family, while, BLASTp finds that this sequence is identical to sequences annotated as the YIP1 family, agreeing with the ProtENN prediction.^2^

Around 40% of Pfam sequences belong to higher-order clans [28]; groups of evolutionarily related families constructed through manual curation involving analysis of structure and function and HMM profile-profile comparison [32, 33]. Clans are often constructed in cases where a single HMM model is not able to capture the full diversity of a large sequence family [32], and individual protein domain sequences can belong to more than one family within a clan. The deep learning models were not given any information about the existence of clan level sequence annotations. Sequence membership in more than one family is addressed in the supplement.

In Supplementary Fig. 2 we report the accuracy of each model at annotating the 55604 held-out test sequences that belong to Pfam clans at both (i) the clan and (ii) the family level. Measuring the accuracy of clan level annotation takes into account the fact that the distinction between different families in the same clan may be less meaningful [31]. Both ProtCNN and ProtENN are able to accurately annotate protein domain sequences at both the clan and the family levels. The error rate of ProtENN is significantly lower for sequence identity in 30-70%; outside this range neither ProtENN or the profile HMMs are significantly better (p < 0.05, McNemar test). All differences, both positive and negative, between ProtCNN and profile HMMs are significant for sequence identity < 70%. At the family level, the neural network models make significantly fewer errors for sequence identities < 80%.

### Performance using a Clustered Split

A key frontier for protein sequence annotation is remote sequence homology detection. The stratified analysis of our random held-out test set suggests that the deep models perform well for distant held-out test sequences, though only 1522 test sequences have identity less than 25% with the train set. To evaluate performance for distant homologs, we use single linkage clustering at 25% identity within each family to split the Pfam-seed data, inspired by [29] (see Methods). This yields a distant held-out test set of 21293 sequences (Supplementary Table 10) that have identity < 25%.^3^ To test the utility of the model identified using the random split, we retain the same model hyperparameters (model architecture, learning rate, etc.) for this significantly harder task. The only variables that were tuned using the held-out clustered dev set were the number of training steps, and the number of ensemble elements for ProtENN (set at 42; larger ensembles were not tested).

A comparison of methods’ overall accuracy appears in Table 2. In Fig. 2 we stratify our analysis of model performance for the clustered split by the percent identity of each held-out test sequence to the closest sequence in the training set. Fig. 2A reports model error rates sliced by sequence identity into 10 bins. This analysis shows that ProtENN performs well across the full range of pairwise sequence identities present. An additional potential performance confounder is family size; large families may be easier to learn, and families with large test sets could skew the reported accuracy. To address the former, we split the held-out clustered test sequence data by total family size into ten bins, each containing ~2100 sequences. Supplementary Fig. 3B shows model error rates for held-out sequences from the clustered split; ProtENN performs well across all family size bins. Finally, in the Supplementary Table 6, we present a table qualitatively similar to Table 2 where the procedure for selecting which sequences from each cluster appear in the test set follows that of [29].

**Table 2:**
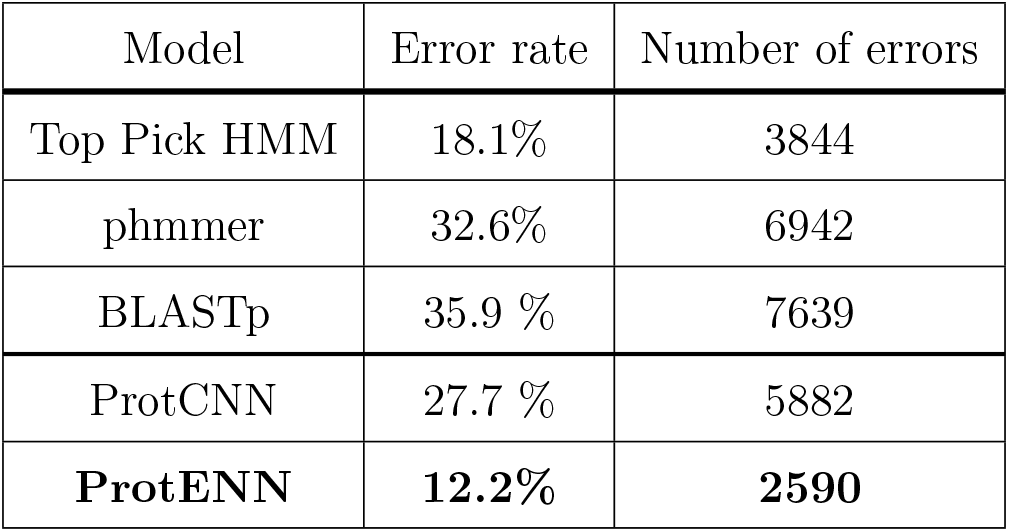
Performance on data split by clustering.

**Figure 2:**
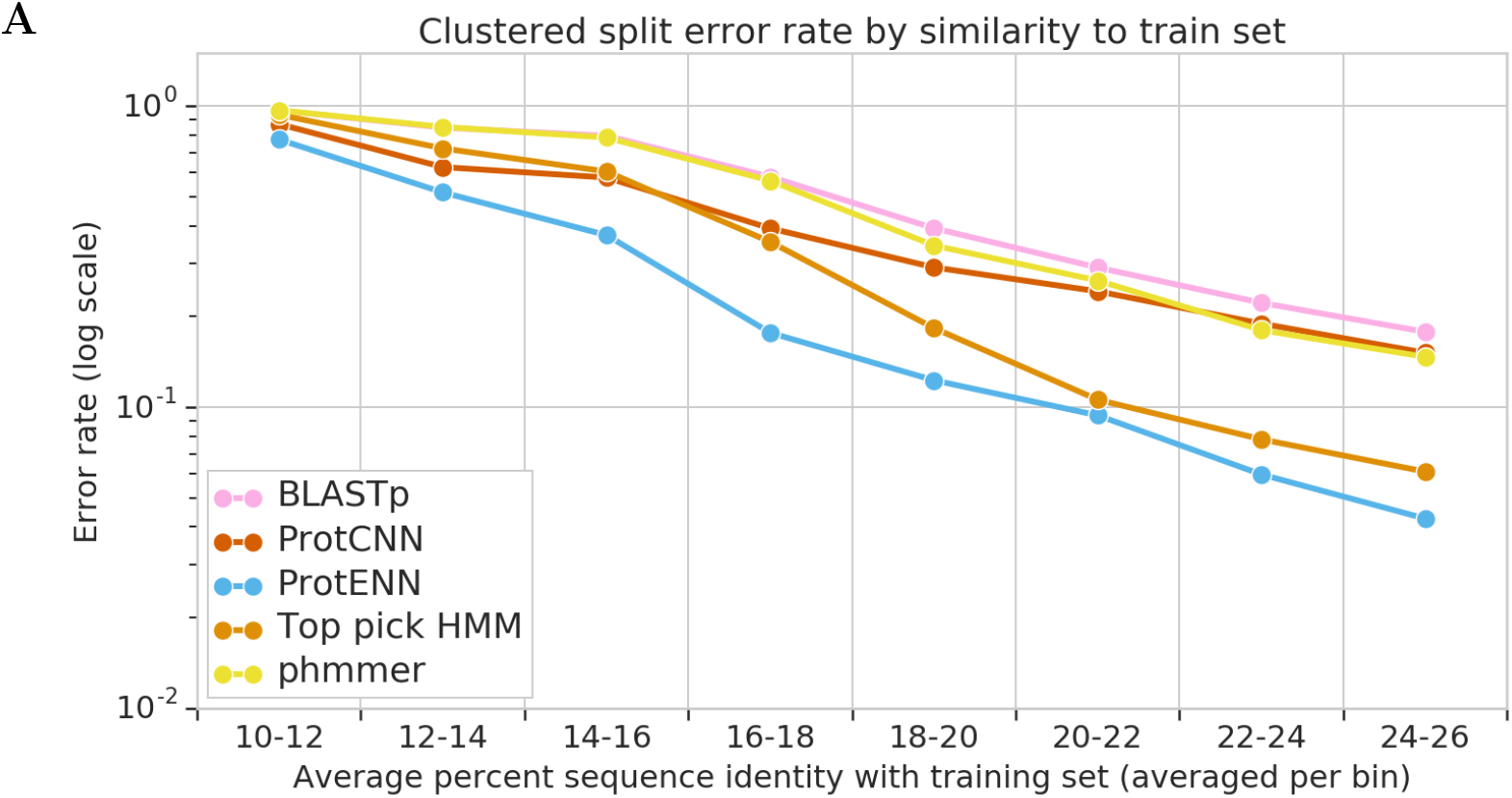
Model performance on the clustered split of Pfam-seed. (A) Held-out test error rate as a function of the maximum percent sequence identity from sequences in the Pfam-seed clustered training set. For each test sequence we use the Pfam-seed family alignment to calculate the percent sequence identity with the most similar train sequence in the same Pfam family (see Methods). Data has been binned by percent sequence identity with the training set; the x-labels describe the bin ranges. ProtENN is significantly better for all bins (p < 0.05, McNemar Test), whereas Top Pick HMM often outperforms ProtCNN. The number of sequences per bin is available in Supplementary Table 5.

### Sequence Annotation for Pfam-full

Pfam-seed contains ~1.34 million curated sequences, and the 17929 profile HMMs built from this data are used to annotate the ~54 million sequences Pfam-full. Like nearest-neighbour methods such as BLASTp, the predictive accuracy of deep learning models typically improves as the amount of well-labelled training data increases. To compare these approaches on a larger dataset, we randomly split each Pfam-full family, assigning 80% of sequences to the train set and 10% each to dev and test sets, and carry out a hyperparameter search to optimize ProtCNN accuracy for this new task. To provide a highly accurate baseline we impute labels via the top BLASTp hit, using the training set as the query database. We do not include profile HMM-based methods (Top Pick HMM and phmmer), because the ground truth data in Pfam full was generated using HMMs.

Our resulting ProtCNN model has an error rate of just 1.26% (~69k errors), lower than the BLASTp error rate of 1.78% (~97k errors). ProtENN, ensembled across 13 ProtCNN models, reduces the error even further to just 0.5% (~25k errors). It is important to stratify our analysis by the similarity of each test sequence to the closest sequence in the training set, to account for sequence similarity between the train and test data. We use the training set as the query database for BLASTp and report the percent sequence identity between members of the highest scoring pair identified by BLASTp for each held-out test sequence. Fig. 3 shows that ProtENN is highly accurate across all bins of held-out test sequences distance from the training data. To analyze the performance for those held-out test sequences that are most distant from the training set, Fig. 3B divides the 90210 held-out test sequences that are most distant from the training sequences into 10 bins, and analyzes model performance for each bin. We find that ProtENN is significantly more accurate for sequences with identity >32% to the training set.

**Figure 3:**
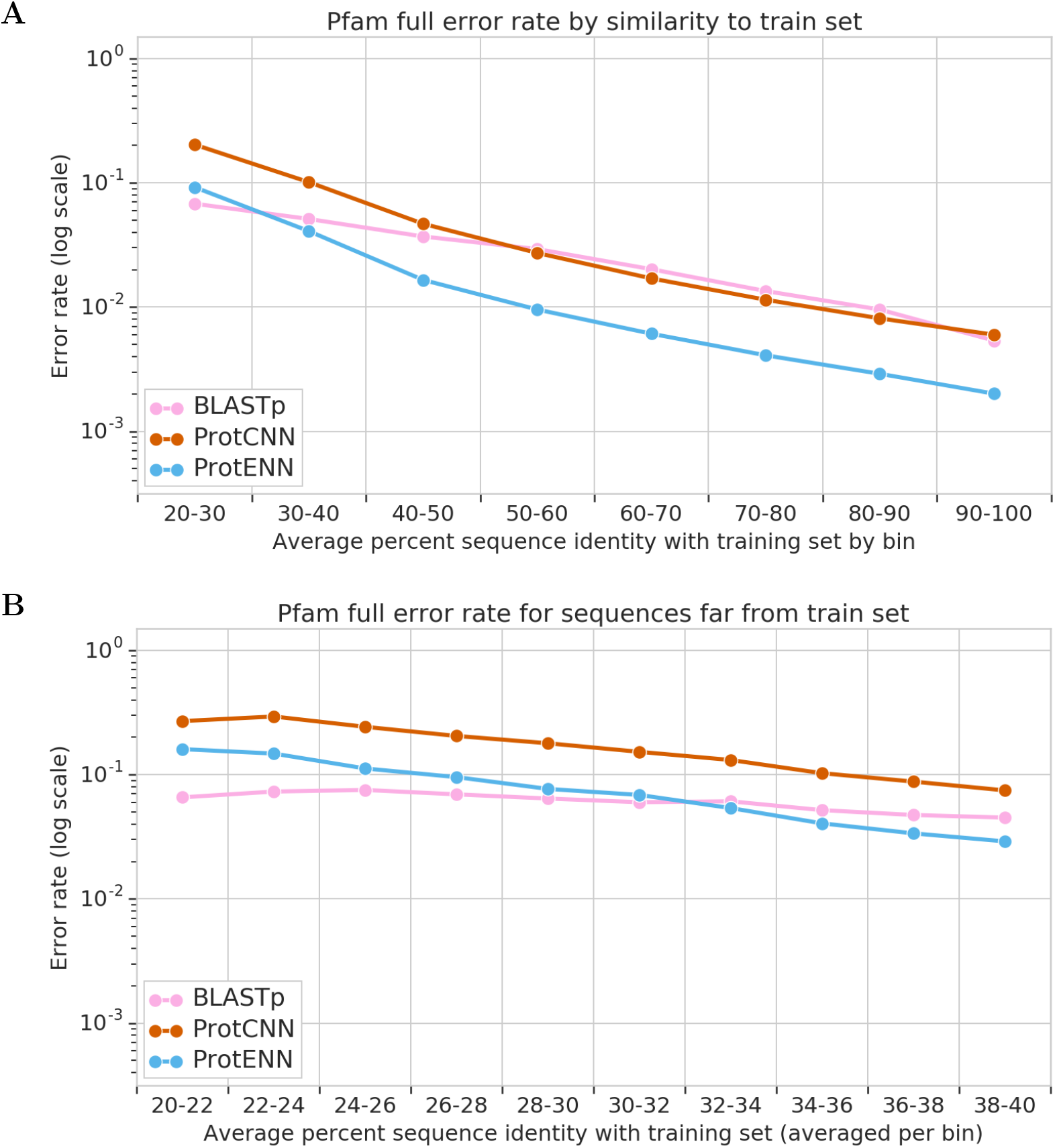
Model performance on the random split of Pfam-full. (A) Held-out test error rate as a function of the percent sequence identity from sequences in the Pfam-full training set. We use the training set as the query database for BLASTp and report the percent sequence identity of the highest scoring pair found by BLASTp for each held-out test sequence. Data has been binned by percent sequence identity with the training set; the x-labels describe the bin ranges. Due to the large test set size the differences between model performance in all bins are statistically significant (p < 0.05, McNemar test). (B) Data for the 90210 sequences with 20-40% sequence identity to the training set were split out, and further subdivided into 10 bins; all differences are statistically significant. The number of sequences per bin in either chart is available in Supplementary Tables 7 and 8.

### Annotation using the learned embeddings

ProtCNN processes an input sequence using two consecutive steps: (1) map the sequence to a 1100-dimensional feature vector (commonly known as an *embedding*) using multiple layers of non-linear transformations, and (2) apply a linear transformation to the embedding to predict a confidence score for each candidate output class. If the overall model performs accurate classification, then sequences from different families will be well-separated in embedding space, since (2) needs to be able to discriminate between the sequences’ embeddings. Using (1) as a general-purpose mapping from sequences to embeddings provides a variety of opportunities beyond the task that the model was initially trained for, including annotation of domains of unknown function and supervised learning on small datasets [19, 26].

We first explore whether the learned embedding space provides an informed metric on unaligned protein sequence data that can be used to accurately annotate sequences that have not been seen during training. We proceed by computing an average embedding for each of the 17929 training families in the seed random split. We then perform nearest-neighbor classification (*Per-Family 1-NN*) for each held-out test sequence using cosine similarity in embedding space with the set of representatives. Note that *Per-Family 1-NN* has the same computational cost as ProtCNN: we remove the final linear transformation layer from ProtCNN and replace it with cosine similarity comparisons to each family’s embedding. More detail is available in the supplement.

In contrast to Per-Family 1-NN, *Per-Instance 1-NN* finds the nearest neighbor for each test sequence among the embeddings of every training sequence (analogous to BLASTp and phmmer). Perhaps surprisingly, Table 3 shows that Per-Family 1-NN is particularly powerful at accurately classifying sequences from small families. Here, performance is analyzed separately for large and small families using the stratification described in the next section. We speculate that Per-Family 1-NN performs better than Per-Instance 1-NN due to noise reduction that results from averaging the embeddings for a family.

**Table 3:**
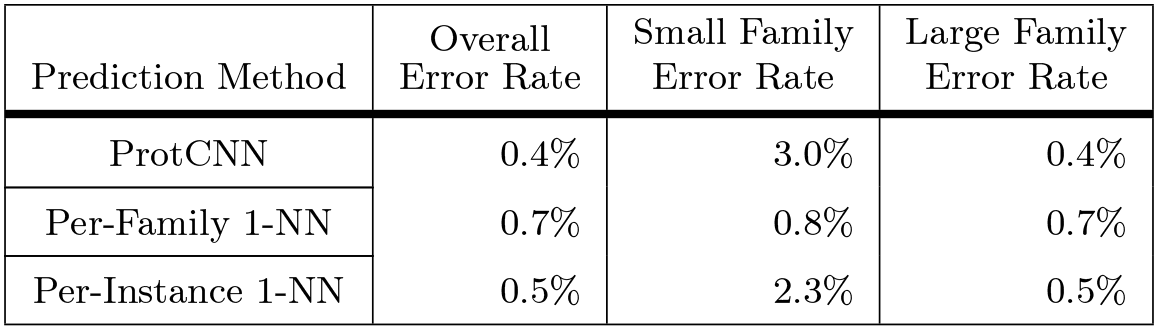
Performance when classifying using nearest neighbors in embedding space.

| Prediction Method | Overall Error Rate | Small Family Error Rate | Large Family Error Rate |
| --- | --- | --- | --- |
| ProtCNN | 0.4% | 3.0% | 0.4% |
| Per-Family 1-NN | 0.7% | 0.8% | 0.7% |
| Per-Instance 1-NN | 0.5% | 2.3% | 0.5% |

In Fig. 4 we apply this approach to the clustered split. We find that *Per-Family 1-NN* considerably improves accuracy vs. ProtCNN, nearly closing the gap with ProtENN. This suggests that the speed-accuracy tradeoff of ProtCNN vs. ProtENN could be avoided using improved machine learning methods, such as [34]. We also demonstrate that computing sequence similarity in terms of embeddings extends well to very remote homologs.

**Figure 4:**
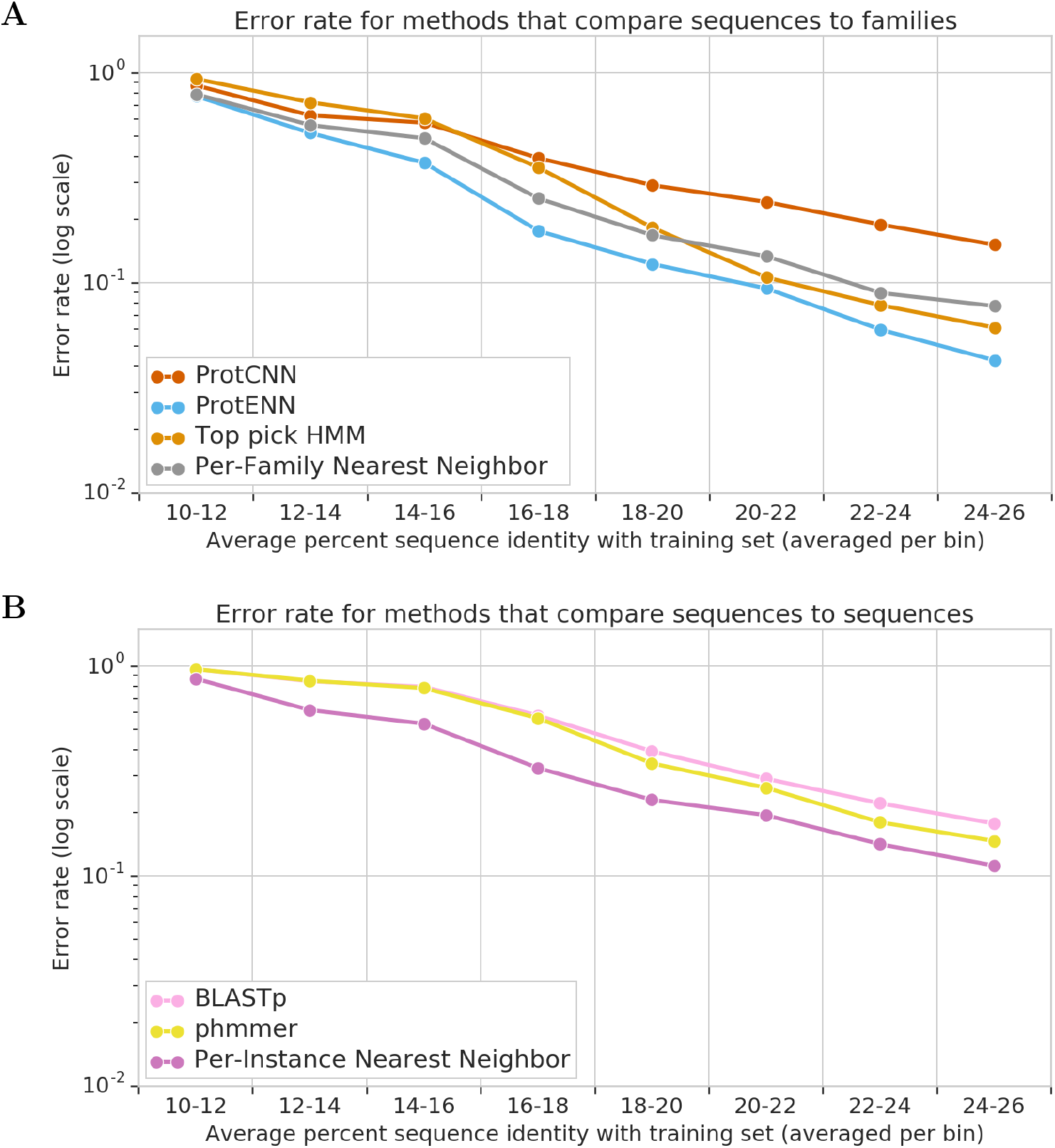
Model performance on the clustered split using the learned embeddings. (A) Error rate of ProtCNN, ProtENN, and per-family nearest neighbors in embedding space. Per-Family 1-NN has the same computational complexity as ProtCNN but is significantly more accurate (p < 0.05, McNemar test) for sequence identity > 12%, with error rate comparable to profile HMMs. This suggests that the speed-accuracy tradeoff between ProtCNN and ProtENN can be side-stepped to yield classifiers that are both faster than ensembles and as accurate. (B) Performance of Per-Instance 1-NN and BLASTp. On this data, sequence similarity using the neural network embeddings enables remote homolog annotation with significantly better accuracy than the pairwise sequence alignment used by BLASTp and phmmer on all bins > 10 (p < .05, McNemar test). The number of sequences per bin in either chart is available in Supplementary Table 5.

### One-Shot Sequence Annotation

Finally, we show that embeddings from ProtCNN can be used to accurately classify sequences from families that the model has not been trained on. This demonstrates further that the deep model is not simply memorizing the training data, and is also motivated by the biologically important question of novel family identification, where each novel family is anchored by a single founder sequence. We proceed by training a ProtCNN model on the subset of Pfam-seed families that have more than 9 training sequences (12361 of 17929 families). The remaining 5568 families consist of 710 families that have a single held out test sequence, and the 4858 smallest families that have no test sequences (because they are so small, see Methods). To simulate the process of introducing new families using a few founder sequences, we exclude these small families from ProtCNN training, but make them available when constructing embeddings for the Per-Family 1-NN approach introduced above.

In Table 4, we compare how various methods perform when given access to only a subset of the available examples for these small families. The number of significant figures on error rates is limited by the size of the test set (710). By construction, ProtCNN has a 100% error rate on the the examples from small families (right column). In contrast, if we include just a single example from each of these families, we can obtain an error rate of 15%, which can be further decreased to 9% if we include two examples. If we had included all available examples for these small classes, the error rate would be 1.3%. It is important to contrast this tradeoff between the number of available training examples and classification performance versus that of HMMER, which is often used by practitioners to grow families using few initial examples [29, 35]. We find that when using only a single founder, HMMER significantly outperforms Per-Family 1-NN, but that this gap is closed if we include two sequences for Per-Family 1-NN. More detail on the few-shot implementations is available in the supplement.

**Table 4:** Contrasting methods’ performance when only a small number of sequences are made available for certain families. This helps demonstrate the extrapolation capabilities of ProtCNN embeddings to regions of sequence space unseen during training, as well shows the these embeddings can help grow families using only a few founder sequences.

| Prediction Method | # Training Examples Used From Each Small Family | Overall Error Rate | Small Family Error Rate |
| --- | --- | --- | --- |
| ProtCNN | 0 | 1.0% | 100.0% |
| Per-Family 1-NN | 1 | 0.8% | 14.9% |
| Per-Family 1-NN | 2 | 0.8% | 9.0% |
| Per-Family 1-NN | all available | 0.7% | 1.3% |
| Top Pick HMM | 1 | 1.4% | 9.3% |
| Top Pick HMM | all available | 1.4% | 1.1% |

### Computational Performance

Protein sequence databases like UniProt contain hundreds of millions of sequences and are growing exponentially [11, 28]. This places a premium on the computational performance of protein sequence analysis tools, motivating efforts dedicated to optimization over the last decades [3, 8, 29, 30, 36, 37]. It is therefore critical to evaluate the computational cost of the deep models to ensure that they aren’t prohibitively expensive. Evaluating the runtime performance of software is delicate. To ensure reproducibility, we use sandboxed instances on Google Cloud Platform: a n1-standard-32 (32-core / 120 GB RAM) instance for CPU-only and a n1-standard-8 (8-core 32GB RAM) + NVIDIA P100 GPU instance for GPU testing. A full set of commands to reproduce our analysis is provided in the supplement.

Table 5 shows the computational performance of ProtCNN, HMMER^4^, and BLAST on our benchmark. ProtCNN on a single CPU processes 9.7 seqs/sec, substantially faster than BLASTp (1.2 seqs/sec) and hmmscan (2.2 seqs/sec) but 2.5x slower than hmmsearch (24.4 seqs/sec). Using the P100 GPU accelerates the inference speed of ProtCNN by a factor of 38, achieving 376.6 seqs/sec. Since both `hmmsearch` and BLASTp run efficiently in parallel, equivalent throughput would require ~15 CPU cores for `hmmsearch` and ~342 cores for BLASTp. Our most accurate model (ProtENN) involves an ensemble of 13 distinct ProtCNN models, implying a throughput of ~29 sequences per second when using a GPU, though distillation [34] would presumably significantly improve this throughput. This demonstrates that the deep learning models presented here can be used with reasonable turn-around times using standard computational resources.

**Table 5:**
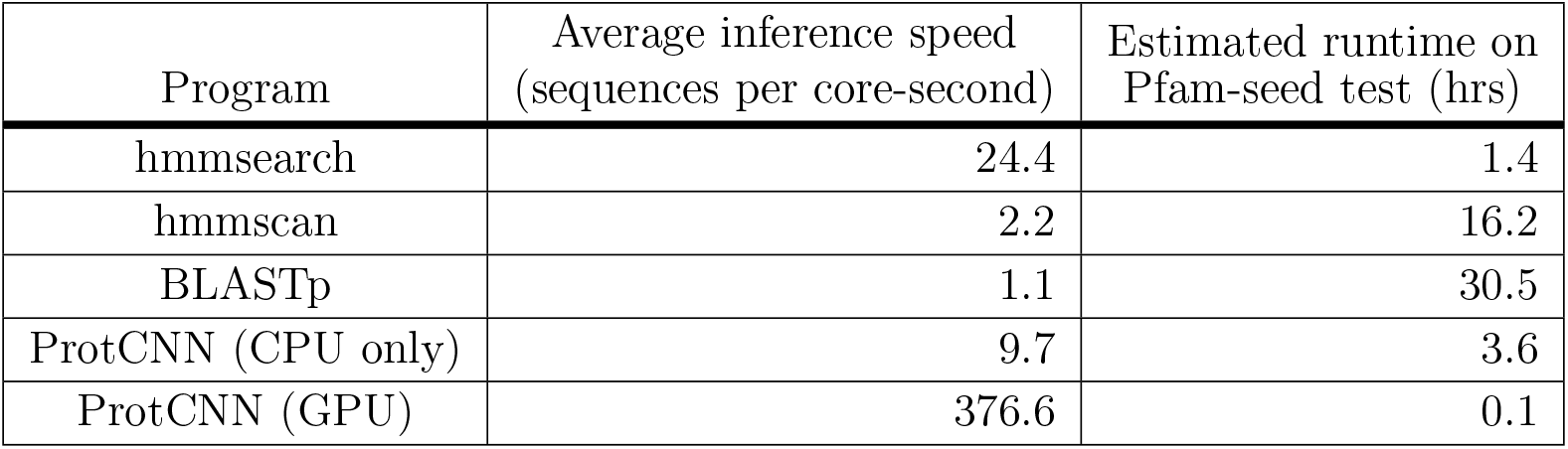
Inference speed of `hmmscan`, `hmmsearch`, and `blastp` run on sandboxed n1-standard-32 (32-core, 120 GB RAM) Google Cloud Platform instances with all data in main memory and using a single core. The ProtCNN model was run in a similar configuration on a n1-standard-8 instance (8-core, 32 Gb RAM) using a single CPU thread for ProtCNN (CPU only), and additionally, one NVIDIA P100 GPU accelerator for ProtCNN (GPU). Additional details, including commands used, are available in the supplement.

### What does ProtCNN learn?

To interrogate what ProtCNN learns about the natural amino acids, we add a 5-dimensional trainable representation between the one-hot amino acid input and the embedding network (see Methods for details), and retrain our ProtCNN model on the same unaligned sequence data from Pfam-full, achieving the same performance. Fig. 5A (left) shows the cosine similarity matrix of the resulting learned embedding, while Fig. 5A (right) shows the BLOSUM62 matrix, created using aligned sequence blocks at roughly 62% identity [38]. The structural similarities between these matrices suggest that ProtCNN has learned known amino acid substitution patterns from the unaligned sequence data.

**Figure 5:**
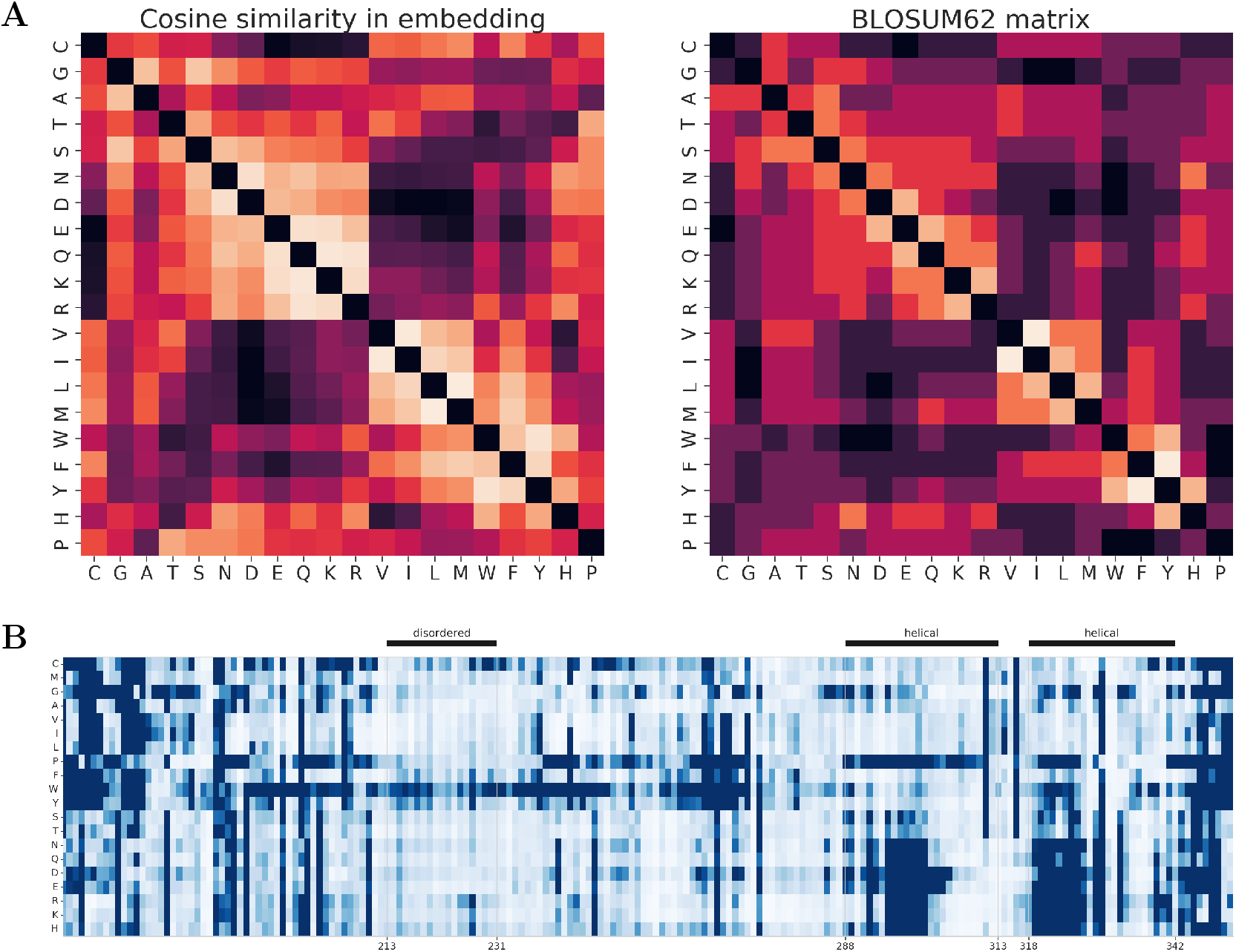
(A) The amino acid embedding extracted from the trained ProtCNN model yields cosine similarities in embedding space that reflect the overall structure of the BLOSUM62 matrix [38]. (B) Predicted change in function for each missense mutation in ATPase domain AT1A1_PIG/161-352 from family PF00122.20. The ProtCNN model (trained using Pfam-full) appropriately predicts that most substitutions in the disordered region are unlikely to change the protein’s function. Substitutions to phenylalanine (P), tyrosine (T) and tryptophan (W) are predicted to have the largest effect on function within the disordered region, in agreement with their known order-promoting properties [39]. The wild type sequence is available in Supplementary Table 9.

We next ask whether ProtCNN can distinguish between variants of the same protein domain sequence with single amino acid substitutions, despite the lack of residue-level supervision during training. To measure the predicted impact of sequence changes, we use a single ProtCNN trained on Pfam-full to calculate the model’s predicted distribution over classes for the original and modified sequences. We then compute the KL-divergence between these two probability distributions to quantify the effect of the substitution on the model prediction. Fig. 5B reports this measure for every possible single amino acid substitution within an ATPase domain sequence. Most substitutions in the disordered region are predicted to have negligible effect, with the exception of mutations to phenylalanine, tyrosine and tryptophan, amino acids that are known to promote order [39]. This ATPase domain also contains two transmembrane helices, within which the order of amino acid (using IUPAC amino acid codes) preference according to ProtCNN is FMLVI YACTS WGQHN KRPED. The suggestion that charged amino acids and proline are avoided within these regions again agrees with existing knowledge [40]. An additional example of saturation mutagenesis prediction is shown in Supplementary Fig. 4.

## Discussion

In this work we compare the performance of ProtCNN (a single deep neural network), ProtENN (an ensemble of ProtCNN models), and methods based on the learned embeddings from ProtCNN against profile HMMs and nearest-neighbor methods on the task of Pfam domain annotation. Given ~1.1 million training examples across 17929 output families of vastly different sizes (Supplementary Fig. 5) our deep models are highly accurate despite having no access to the alignments used by the profile HMMs. These results present a significant advance over previous efforts applying deep learning methods in terms of the number of families, and the number of training sequences per family. On randomly split data, ProtCNN achieves an error rate of only 0.495% and ProtENN has an error rate of 0.159%, compared to 1.14%, 1.531%, and 1.654% for HMMER, phmmer, and BLASTp, respectively. The difference in performance is significant across a wide range of similarities between test sequences and the training set. To further evaluate the models’ performance for even more remote homologs, we consider a clustered train-test split. Here, we find that ProtENN, with an overall error rate of 12.2%, significantly outperforms HMMer, which has an error rate of 18.1%, while ProtCNN has an error rate of 27.7%. Methods that use the embeddings produced by ProtCNN can accurately cluster sequences from unknown families, providing further confidence that the models learn an informative representation of protein domain sequence data.

Importantly, we stratify model performance by percent sequence identity with the training data for every data split we construct, which serves to avoid overestimating the generalization capabilities of the model. In addition to stratifying performance for a random split, we construct a clustered split in which all held-out test sequences are guaranteed to be far from the train set. The community has embraced the second evaluation approach, but we maintain that the former is at least as important. If future users of such a machine learning system will issue prediction requests for sequences that are drawn from a distribution similar to the existing data, the random split helps us evaluate how useful the system will be to these researchers. Furthermore, performing a stratified analysis of the randomly-split data reveals how performance varies with sequence identity without introducing systematic skew in the training data due to clustering. On the other hand, if users will mostly issue queries for very remote sequences, then evaluating models in terms of the clustered split is important.

When choosing between ProtCNN and ProtENN there is a clear speed-accuracy tradeoff. However, the ability of the embedding-based approaches to improve the single model accuracy, without adding computational overhead, suggests that this tradeoff is not necessary (Fig. 4A). Application of more sophisticated machine learning methods have the potential to lead to further performance gains. Such approaches could involve, for example, distilling the ensembled CNN models into a single model [34].

The representation of protein sequence space learned by ProtCNN is also used in one-shot learning to classify sequences from unseen protein families. This suggests an iterative approach to novel family construction, inspired by current methods such as PSI-BLAST [3], jackhmmer [29, 35] and hhblits [4]. A single founder sequence is used to find additional family members, which are then used to update the average embedding for this putative new family and so forth. Our results suggest that a deep model trained on an existing corpus of data (here the training sequences from large Pfam seeds) could accurately build a new family from a single sequence even in the presence of additional sequences from families that the model was not trained on. Future work will test this approach beyond the confines of the benchmark Pfam-seed dataset.

Our deep models achieve extremely high accuracy without prior knowledge of protein sequence data encoded through substitution matrices, sequence alignment or hand-curated scoring functions. The embedding network in each ProtCNN model maps an input sequence to a single vector representation that alone can be used for accurate family classification, pairwise sequence comparison or other downstream analysis. This differs substantially from approaches such as BLASTp, phmmer and HMMER that perform annotation using explicit alignment. We note that simpler models provide useful attribution of model decision making, and we anticipate that similar insights will emerge from work that improves the interpretation and understanding of deep learning models [41–43].

In this work, we focus on protein domain sequence annotation to provide a benchmark with broad coverage that enables comparison with the state of the art profile HMMs provided by Pfam 32.0. Though carefully curated, at least 25% of sequences have no experimentally validated function [28], and additional functional characterization of protein sequences would greatly improve model quality. Furthermore, the model training protocol that we describe can be applied to any set of labelled protein sequence data and our results suggest that deep learning models can rapidly and efficiently annotate novel protein sequences. Such approaches have the potential to unlock novel molecular diversity for both therapeutic and biotechnology applications.

## Methods

### Deep Learning Models

We use unaligned sequence data to train deep models that learn the distribution across protein families through joint optimization of a softmax regression loss function. Fig. 6 depicts the *input*, *embedding* and *prediction* networks that make up each deep learning model. The input and prediction networks have the same functional form for all models. The input network maps a sequence of *L* amino acids to an *L* × 20 binary array, where each column is a one-hot amino acid representation (Fig. 6A). Sequences are padded to the length of the longest sequence in the batch with all-zero vectors on the right. The embedding network maps the *L* × 20 array containing the one-hot amino acid representation of the sequence to an *L* × *F* array that contains an embedding for each sequence residue (Fig. 6B). *F* is a tunable hyperparameter, which we set to 1100 for ProtCNN. For residues outside the set of the 20 natural amino acids, we use a column of zeros. All processing in the subsequent embedding network is designed such that it is invariant to the padding that was introduced for a given sequence. Additional detail about network architectures are available in the supplement, and neural network hyperparameters that were tuned using the development set are provided in Supplementary Tables 11, 12, 13 and 14.

**Figure 6:**
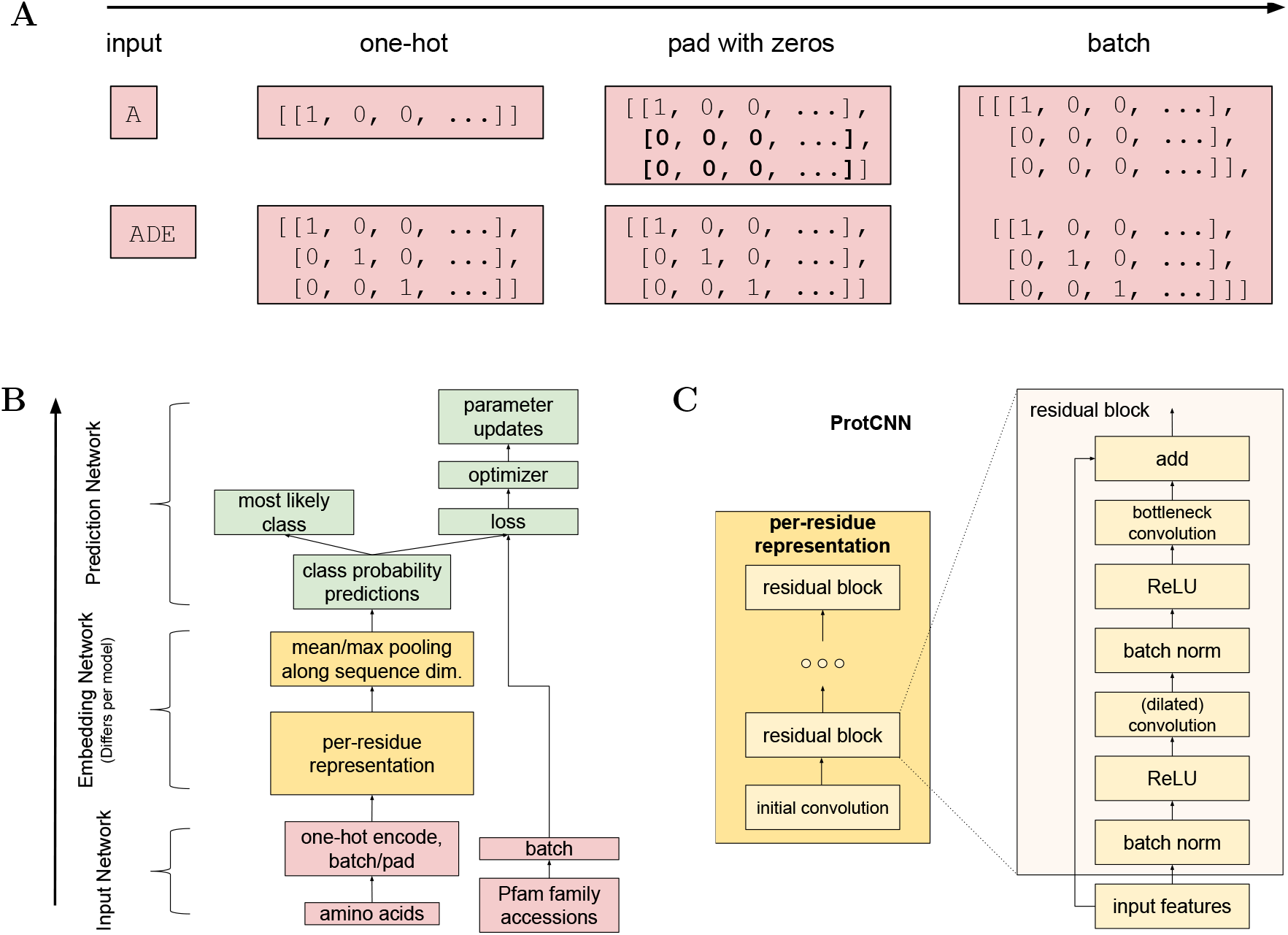
(A) Comparison of the within Pfam family distance calculated using the formula given in the text with the BLASTp percent sequence identity for each of the 126171 held-out test sequences of the Pfam-seed dataset. (B) Number of test-set sequences for each family in the random and clustered splits. While the clustering process is desirable because it ensures separation between train and test data, it introduces a distribution over families in the test data that is significantly different than the overall distribution in Pfam.

For the embedding network, ProtCNN uses convolutional residual networks (ResNets [44], a variant of convolutional neural networks that in practice train quickly and are stable, even with many layers [44]). Fig. 6C depicts the ResNet architecture, which includes dilated convolutions [45]. The ProtCNN networks are translationally invariant, an advantage for working with unaligned protein sequence data. An n-dilated 1d-convolution takes standard convolution operations over every nth element in a sequence, allowing local and global information to be combined without greatly increasing the number of model parameters. An important composite hyperparameter is the *receptive field size* of each per-residue feature, which describes the length of the subsequence that affects its value. Using dilated convolutions enables larger receptive field sizes without an explosion in the number of model parameters. For this benchmark setup, we find that larger receptive fields generally correspond to higher accuracy (Supplementary Fig. 1). To our knowledge, this is the first application of dilated convolutions to protein sequence annotation.

The *L* × *F* array is pooled along the length of the sequence to produce an *F*-dimensional embedding by taking the maximum over each row. The prediction network maps the output of the embedding network *F* to a distribution over labels using a multi-class logistic regression model, where the vector of probabilities is obtained as SoftMax(*Wf* + *b*), where *f* ∈ *F* and *W* and *b* are learned weights and biases. The model prediction is the most likely label under this distribution. At train time, the log-likelihood and its gradient with respect to the parameters of the prediction and embedding networks are computed using standard forward and back propagation.

ProtCNN is orders of magnitude faster at making predictions than BLASTp; the basic numerical operations required can be parallelized both along the length of the sequence and across multiple sequences, and can be accelerated by hardware. In addition to the CNN models we also trained a recurrent neural network (RNN) with single-layer bidirectional LSTM [46], which achieved accuracy of 0.982 on the Pfam-seed dataset. Replicate Deep CNN models trained on different orderings of the same data with different random parameter initializations make distinct errors, leading to ProtENN, an ensemble of ProtCNN models.

**Figure 6:**
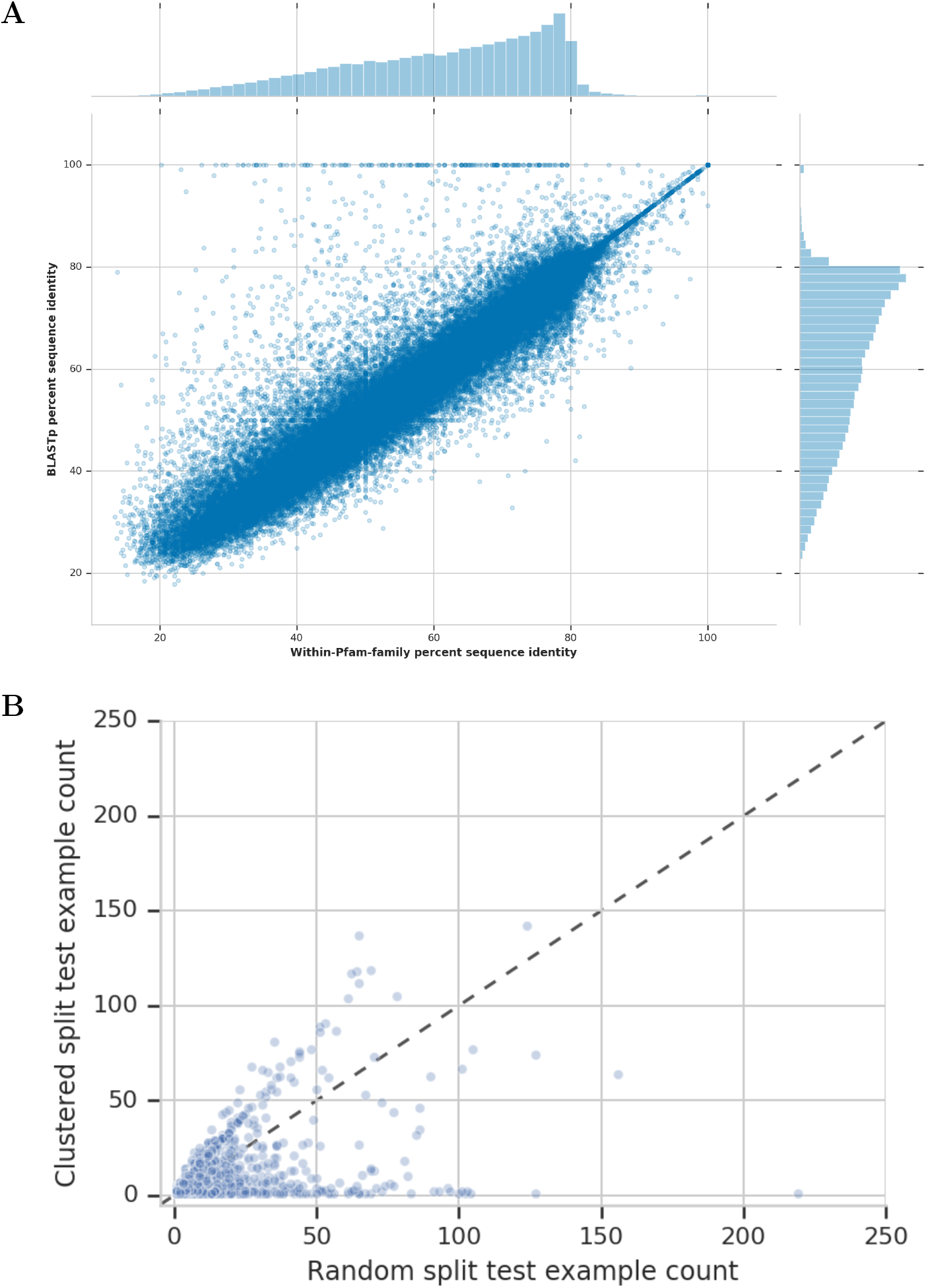
Architecture descriptions of neural networks. (A) Adding padding to a sequence for batching. (B) The model architecture shared by all neural networks includes the Input (red), Embedding (yellow), and Prediction (green) Networks. (C) ResNet architecture used by the ProtCNN models.

Overall those ProtCNN models that perform best tend to have the largest memory footprint, to some extent irrespective of how that memory footprint is achieved. Increasing the number of model parameters via the number of filters, the kernel size and/or the number of ResNet blocks, and increasing the training batch size can all produce performance improvements. Fundamentally, the memory footprint of the models we trained was limited by the amount of memory available on a single GPU, necessitating trade offs among these different factors. Additional computational resources can overcome this memory limitation: we didn’t explore TPUs [47], multiple GPUs or CPUs, all of which could result in better models. This suggests that there is room for future machine learning developments on this task.

### Benchmark Dataset for Random Split

To benchmark different models at unaligned protein domain sequence annotation we turn to the highly curated Protein families (Pfam) database [28, 48]. The 17929 families of the Pfam 32.0 release are labelled using HMMs that provide broad coverage of the known protein universe; 77.2% of the ~137 million sequences in UniprotKB have at least one Pfam family annotation, including 74.5% of proteins from reference proteomes [11, 28]. Many domains have functional annotations, although at least 22% of Pfam domains have unknown function [28]. The HMM for each Pfam family is built from a manually curated family seed alignment, containing between 1 and 4545 protein domain sequences of length 4-2037 amino acids (Supplementary Fig. 5).

We split each Pfam family with at least 10 seed sequences randomly into disjoint dev^5^ (10%, rounding down to the nearest integer) and test (10%) sets, allocating remaining sequences to the training set. Of the 17929 Pfam-seed families, 4858 families have < 10 seed sequences and are only present in the train set. This results in held-out test sequences for 13071 families, where 2819 families have exactly one sequence in each of the test and dev sets. Including these additional families in the train set makes the task harder because there are more ways each test sequence can be misclassified. Note that we do not expect the HMM-based approach to achieve 100% accuracy because the training data is a subset of the seed data set used in Pfam.

For reproducibility, we provide the split Pfam seed dataset for download,^6^ and at Kaggle, together with an interactive Jupyter notebook.^7^ For the profile HMMs, we retain the alignment information from the whole Pfam-seed for all splits to avoid any artifacts introduced by realignment, and enable optimal performance. During training this provides the HMM with information about the held-out test sequences used to measure performance, meaning that the reported accuracy should be taken as an upper bound. In contrast all alignment information is removed from the data for our deep learning models and for the other baselines.

### Definition of sequence identity

Inspired by the method of [49], we use the Pfam-seed family alignments to compute the similarity, measured as percent sequence identity, between every held-out test sequence and the training sequences within the same family. For each pair containing a single test and train sequence we do not realign, but instead retain their length *L* alignment from the family multiple sequence alignment. Following [49], for two sequences of *n*_1_ and *n*_2_ residues, if the *L* aligned sequence position pairs consist in *c*_ident_ matches, *c*_ident_ mismatches and *c*_ident_ cases where either or both sequence contain a gap character such that *L* = *c*_ident_ + *c*_mismat_ + *c*_indel_, then the pairwise sequence identity is defined as:

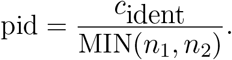

For the Pfam-full data (see Fig. 3) we instead use BLASTp to calculate a measure of sequence identity. We use the Pfam-full training set as the query database for BLASTp and report the percent sequence identity of the highest scoring pair found by BLASTp for each held-out test sequence. This method measures similarity across all Pfam families, in contrast to the method described above, which computes the distance between the train and test sets within each Pfam family. Supplementary Fig. 6 compares these two metrics across the 126171 held-out sequences of the randomly split Pfam-seed data.

### Benchmark Dataset for Clustered Split

Fig 1 shows that our randomly split train and test sets contain some examples with high sequence identity. A model tuned on such data might be suspected to perform poorly at *remote homology detection*, [29]. To address this, we apply single-linkage hierarchical agglomerative clustering to each family to build clustered train, dev and test sequence sets that are guaranteed to be distant in sequence space from each other, as enforced by the protocol developed in [29]. Note that our clustering protocol follows that of [29], but we evaluate models in terms of a different prediction task. We consider multi-class classification, whereas [29] considers a set of per-family binary detection problems.

For a given family, we split the sequences into set1 and set2 using the following steps: (1) Construct the matrix of pairwise distances among family members, where distance is measured in terms of sequence identity according to the Pfam-seed family alignment. (2) Run single-linkage hierarchical agglomerative clustering and process the resulting clustering tree to yield a set of clusters where each element of a given cluster is guaranteed to have at most α sequence identity with the nearest element of any other cluster. (3) Sort the clusters by size and add clusters to set1, until its size exceeds a fraction Δ of the overall family size. We first use this procedure to split the data into training and non-training sets. We set α = 0.25 and Δ = 0.5. We then recluster the non-training data at a threshold of α = 0.7, and follow the procedure above to split the non-training data into a dev and test set. For the results presented in this paper, we use the clustered dev data to tune the number of training iterations and the number of ensemble elements, and make no changes to the model hyperparameters from those identified using the random split.

Overall, our approach follows that of [29] with four main modifications. The first is that we place multiple clusters in set1, rather than just the largest. This avoids putting very few examples in the training set for families where the clustering produces a large number of small clusters, while maintaining the property that the train, dev, and test sets are well-separated. Second, our formulation uses some of the non-train sequences for a dev set to make sure that the number of training steps and ensemble elements are not chosen using the held-out test data. Third, if a family can not be split at sequence identity α, we place the entire family in the training set. This differs from [29], which completely discards families that can not be split. When clustering non-train data to split into dev and test, we similarly place the entire family in the dev set if it can not be split. The fourth is that when we re-cluster the non-training data to produce dev and test sets, instead of selecting single sequences from each cluster, we include all elements of each cluster. We find that though the fourth decision simplifies our setup, following more closely [29] yields qualitatively similar results (Supplementary Table 6).

Finally, when considering aggregate accuracy metrics it is important to consider the test distribution under which this metric is computed. The randomly-split test data (especially for Pfam-full) has a natural distribution over families defined by the distribution over families in Pfam. However, in the clustered data the distribution over families is a complex consequence of the clustering process. In Supplementary Fig. 6B, we find that many families are represented very differently in the randomly and clustered data.

An additional potential performance confounder is family size. To address this issue, we split the held-out test sequence data for the Pfam-seed random and clustered splits and also for the Pfam-full random split by total family size into ten bins. Supplementary Fig. 3 shows the model error rate for held-out sequences from each data split. These results show that ProtENN performs well across all family size bins.

### Baseline Classifiers

#### phmmer

We take the set of unaligned training sequences as a sequence database, and using the `phmmer` function from HMMER 3.1b [29] we query each test sequence against this database to find the closest match. Those test sequences that return hits above the default phmmer reporting threshold are then annotated with the label of the training sequence hit with the highest bit score. Out of the 126171 sequences in the test set, 42 did not return a hit using this approach. All training sequences that are not reported as hits by the phmmer function are assumed to have a zero bit score match to that query sequence.

Our strategy of working with the Pfam-seed sequence set circumvents the computationally intensive process of evaluating phmmer on the full set of ~54.5 million Pfam sequences.

#### k-mer

An alignment-free approach is provided by a k-mer (or n-gram) based model, where each sequence is represented by the set of k-mers that it contains. We train a multi-class logistic regression model on vectors of k-mer counts using the same stochastic gradient descent procedure as used by our deep models (Supplementary Table 15).

#### BLASTp

BLASTp [30] is one of the most well known algorithms for searching for similar sequences, and among the current state of the art. It uses an alignment to rank sequences according to their similarity to a target sequence, and from this, a user can impute functional annotation by ascribing known functions of similar sequences. We use BLASTp as a 1-nearest neighbor algorithm by first using `makeblastdb` (version 2.7.1+) with the training data. We then query sequences from that database using blastp -query, taking only the top hit. This implementation returns no hit for 259 (0.21%) of the 126171 sequences in the Pfam-seed test set.

#### Top Pick HMM Implementation

Profile HMMs are widely regarded as a state of the art modeling technique for protein sequence annotation. We used `hmmbuild` from HMMER 3.1b to construct a profile HMM from the aligned train sequences for each family in Pfam 32.0. We implement a simple top pick HMM strategy to avoid any handicap from the filters built into HMMER 3.1b. To further obtain the best possible profile HMM performance, we retain the alignment from the entire Pfam-seed, avoiding dependence on any particular realignment method. We then use the `hmmsearch` function from HMMER 3.1b to search all 17929 profiles against the set of unaligned test sequences using the default parameter settings.

The scores for each hit are recorded, and the profile with the highest score called as the HMM prediction for that test sequence. The HMMs yield no prediction for 445 sequences (0.35%) of the test set, increasing the number of errors to 2229 over the value given in Table 1. To ensure the HMMER 3.1b heuristics did not hamper performance, we manually turn them off to the extent that at least one hit is reported for each test sequence, and take the top scoring hit. To implement this, for those test sequences with no profile hit after this first pass, we employ a second `hmmsearch` pass using the `--max` option, which turns off all filters and runs full Forward/Backward inference on every target to increase the sensitivity of the search at a significant cost in speed [50].

In experiments that retain the HMMER 3.1b filters for `hmmsearch`, we found that 8.5% of test sequences returned multiple hits above the family specific gathering thresholds that are used by Pfam to regulate family membership. Reporting these results would have resulted in a lower precision score for HMMER than for the deep models, which is one of the reasons we have chosen instead to remove the statistical filter and report the top hit (Supplementary Fig. 5).

The positive results obtained in the absence of rigorous statistical filters likely reflect the fact that we are working with sequences that were originally classified by Pfam, and so passed the rigorous statistical thresholds set for inclusion in a Pfam family. Those sequences that did not pass these filters, and hence were not included in any Pfam family, may well have posed a more significant challenge to our implementation. For this reason we do not recommend that this HMM implementation is used in settings other than working with these benchmark datasets. For Pfam-full, we do not use the HMMs as a baseline because these models were used to label the data, so may achieve 100% accuracy by default. The Pfam-full dataset has 17772 families overall, and our test and dev sets contain sequences from 16755 families.

### Computational Performance

Table 5 reports the number of sequences processed per core-second, computed using the runtime to process 10% of the seed test fasta sequences. We limited each program to a single CPU core to focus on computational efficiency rather than the effectiveness of shared memory parallelization. To minimize the cost of input/output (IO), all data files were held in RAM (see Supplementary Materials for details). We ran inference for ProtCNN both with and without a GPU accelerator. The GPU configuration represents a common inference environment for deep learning models, while the CPU-only configuration allows direct comparison with BLASTp and HMMER. We made a good faith effort to build and run all programs efficiently in this environment; additional details, including command lines for benchmarking, are available in the Supplementary Materials. Note that GPU-accelerated versions of BLASTp [51] and HMMER [52] were not evaluated and may have significantly higher throughput than the CPU-only versions considered here.

## Acknowledgements

We thank Jamie Smith for countless conversations and guidance throughout this project; Eli Bixby for an implementation of ragged tensor processing that sped up our ProtCNN implementation substantially on GPU; Cory McClean, Babak Alipanahi, and Steven Kearnes for extensive proofreading and feedback.

## Footnotes

1 Available for download at https://console.cloud.google.com/storage/browser/brain-genomics-public/research/proteins/pfam/random_split, interactive notebook at https://www.kaggle.com/googleai/pfam-seed-random-split

2 DUF1282 will soon be merged with YIP1 in Pfam.

3 https://console.cloud.google.com/storage/browser/brain-genomics-public/research/proteins/pfam/clustered_split

4 We benchmark two methods of comparing HMMs to sequences, hmmsearch and hmmscan, which are both provided by the software package HMMER. More detail can be found in the Computational Performance section in the supplement.

5 A dev (development) set is a set used to tune hyperparameters that is separate from the test set to avoid overfitting on test data.

6 https://console.cloud.google.com/storage/browser/brain-genomics-public/research/

7 https://www.kaggle.com/googleai/pfam-seed-random-splitproteins/pfam/random_split

## References

[1] Martin Steinegger and Johannes Söding. Mmseqs2 enables sensitive protein sequence searching for the analysis of massive data sets. Nature biotechnology, 35(11):1026, 2017.

[2] Martin Steinegger and Johannes Söding. Clustering huge protein sequence sets in linear time. Nature communications, 9(1):2542, 2018.

[3] Stephen F Altschul, Thomas L Madden, Alejandro A Schäffer, Jinghui Zhang, Zheng Zhang, Webb Miller, and David J Lipman. Gapped blast and psi-blast: a new generation of protein database search programs. Nucleic acids research, 25(17):3389–3402, 1997.

[4] Johannes Söding. Protein homology detection by hmm–hmm comparison. Bioinformatics, 21(7):951–960, 2004.

[5] Robert D Finn, Jody Clements, and Sean R Eddy. Hmmer web server: interactive sequence similarity searching. Nucleic acids research, 39(suppl_2):W29–W37, 2011.

[6] Morgan N Price, Kelly M Wetmore, R Jordan Waters, Mark Callaghan, Jayashree Ray, Hualan Liu, Jennifer V Kuehl, Ryan A Melnyk, Jacob S Lamson, Yumi Suh, et al. Mutant phenotypes for thousands of bacterial genes of unknown function. Nature, page 1, 2018.

[7] Yi-Chien Chang, Zhenjun Hu, John Rachlin, Brian P Anton, Simon Kasif, Richard J Roberts, and Martin Steffen. Combrex-db: an experiment centered database of protein function: knowledge, predictions and knowledge gaps. Nucleic acids research, 44(D1): D330–D335, 2015.

[8] Noah M Daniels, Andrew Gallant, Jian Peng, Lenore J Cowen, Michael Baym, and Bonnie Berger. Compressive genomics for protein databases. Bioinformatics, 29(13): i283–i290, 2013.

[9] Robert D Finn, Alex Bateman, Jody Clements, Penelope Coggill, Ruth Y Eberhardt, Sean R Eddy, Andreas Heger, Kirstie Hetherington, Liisa Holm, Jaina Mistry, et al. Pfam: the protein families database. Nucleic acids research, 42(D1):D222–D230, 2013.

[10] Yann LeCun, Yoshua Bengio, and Geoffrey Hinton. Deep learning. nature, 521(7553): 436, 2015.

[11] UniProt Consortium. Uniprot: the universal protein knowledgebase. Nucleic acids research, 45(D1):D158–D169, 2016.

[12] Jie Hou, Badri Adhikari, and Jianlin Cheng. Deepsf: deep convolutional neural network for mapping protein sequences to folds. Bioinformatics, 34(8):1295–1303, 2017.

[13] Maxat Kulmanov, Mohammed Asif Khan, and Robert Hoehndorf. Deepgo: predicting protein functions from sequence and interactions using a deep ontology-aware classifier. Bioinformatics, 34(4):660–668, 2017.

[14] Renzhi Cao, Colton Freitas, Leong Chan, Miao Sun, Haiqing Jiang, and Zhangxin Chen. Prolango: protein function prediction using neural machine translation based on a recurrent neural network. Molecules, 22(10):1732, 2017.

[15] Yu Li, Sheng Wang, Ramzan Umarov, Bingqing Xie, Ming Fan, Lihua Li, and Xin Gao. Deepre: sequence-based enzyme ec number prediction by deep learning. Bioinformatics, 34(5):760–769, 2017.

[16] Balázs Szalkai and Vince Grolmusz. Near perfect protein multi-label classification with deep neural networks. Methods, 132:50–56, 2018.

[17] Balázs Szalkai and Vince Grolmusz. Seclaf: a webserver and deep neural network design tool for hierarchical biological sequence classification. Bioinformatics, 34(14):2487–2489, 2018.

[18] Zhenzhen Zou, Shuye Tian, Xin Gao, and Yu Li. mldeepre: Multi-functional enzyme function prediction with hierarchical multi-label deep learning. Frontiers in genetics, 9, 2018.

[19] Ariel S Schwartz, Gregory J Hannum, Zach R Dwiel, Michael E Smoot, Ana R Grant, Jason M Knight, Scott A Becker, Jonathan R Eads, Matthew C LaFave, Harini Eavani, et al. Deep semantic protein representation for annotation, discovery, and engineering. BioRxiv, page 365965, 2018.

[20] Da Zhang and Mansur R Kabuka. Protein family classification with multi-layer graph convolutional networks. In 2018 IEEE International Conference on Bioinformatics and Biomedicine (BIBM), pages 2390–2393. IEEE, 2018.

[21] Xueliang Liu. Deep recurrent neural network for protein function prediction from sequence. arXiv preprint arXiv:1701.08318, 2017.

[22] Babak Alipanahi, Andrew Delong, Matthew T Weirauch, and Brendan J Frey. Predicting the sequence specificities of dna-and rna-binding proteins by deep learning. Nature biotechnology, 33(8):831, 2015.

[23] Sai Zhang, Jingtian Zhou, Hailin Hu, Haipeng Gong, Ligong Chen, Chao Cheng, and Jianyang Zeng. A deep learning framework for modeling structural features of rna-binding protein targets. Nucleic acids research, 44(4):e32–e32, 2015.

[24] Ehsaneddin Asgari and Mohammad RK Mofrad. Continuous distributed representation of biological sequences for deep proteomics and genomics. PloS one, 10(11):e0141287, 2015.

[25] Sam Sinai, Eric Kelsic, George M Church, and Martin A Nowak. Variational auto-encoding of protein sequences. arXiv preprint arXiv:1712.03346, 2017.

[26] Ethan C Alley, Grigory Khimulya, Surojit Biswas, Mohammed AlQuraishi, and George M Church. Unified rational protein engineering with sequence-only deep representation learning. bioRxiv, page 589333, 2019.

[27] Alexander Rives, Siddharth Goyal, Joshua Meier, Demi Guo, Myle Ott, C Lawrence Zitnick, Jerry Ma, and Rob Fergus. Biological structure and function emerge from scaling unsupervised learning to 250 million protein sequences. bioRxiv, page 622803, 2019.

[28] Sara El-Gebali, Jaina Mistry, Alex Bateman, Sean R Eddy, Aurélien Luciani, Simon C Potter, Matloob Qureshi, Lorna J Richardson, Gustavo A Salazar, Alfredo Smart, et al. The pfam protein families database in 2019. Nucleic acids research, 47(D1):D427–D432, 2018.

[29] Sean R Eddy. Accelerated profile hmm searches. PLoS computational biology, 7(10): e1002195, 2011.

[30] Stephen F Altschul, Warren Gish, Webb Miller, Eugene W Myers, and David J Lipman. Basic local alignment search tool. Journal of molecular biology, 215(3):403–410, 1990.

[31] Alex Bateman. What are these new families with 2, 3, 4 endings?, January 2012. URL https://xfam.wordpress.com/2012/01/19/what-are-these-new-families-with-_2-_3-_4-endings/. [Online; posted 19-January-2012].

[32] Robert D Finn, Jaina Mistry, Benjamin Schuster-Böckler, Sam Griffiths-Jones, Volker Hollich, Timo Lassmann, Simon Moxon, Mhairi Marshall, Ajay Khanna, Richard Durbin, et al. Pfam: clans, web tools and services. Nucleic acids research, 34(suppl_1): D247–D251, 2006.

[33] Robert D Finn, Penelope Coggill, Ruth Y Eberhardt, Sean R Eddy, Jaina Mistry, Alex L Mitchell, Simon C Potter, Marco Punta, Matloob Qureshi, Amaia Sangrador-Vegas, et al. The pfam protein families database: towards a more sustainable future. Nucleic acids research, 44(D1):D279–D285, 2015.

[34] Geoffrey Hinton, Oriol Vinyals, and Jeff Dean. Distilling the knowledge in a neural network. arXiv preprint arXiv:1503.02531, 2015.

[35] L Steven Johnson, Sean R Eddy, and Elon Portugaly. Hidden markov model speed heuristic and iterative hmm search procedure. BMC bioinformatics, 11(1):431, 2010.

[36] Yongan Zhao, Haixu Tang, and Yuzhen Ye. Rapsearch2: a fast and memory-efficient protein similarity search tool for next-generation sequencing data. Bioinformatics, 28 (1):125–126, 2011.

[37] Benjamin Buchfink, Chao Xie, and Daniel H Huson. Fast and sensitive protein alignment using diamond. Nature methods, 12(1):59, 2015.

[38] Steven Henikoff and Jorja G Henikoff. Amino acid substitution matrices from protein blocks. Proceedings of the National Academy of Sciences, 89(22):10915–10919, 1992.

[39] Andrew Campen, Ryan M Williams, Celeste J Brown, Jingwei Meng, Vladimir N Uversky, and A Keith Dunker. Top-idp-scale: a new amino acid scale measuring propensity for intrinsic disorder. Protein and peptide letters, 15(9):956–963, 2008.

[40] C Nick Pace and J Martin Scholtz. A helix propensity scale based on experimental studies of peptides and proteins. Biophysical journal, 75(1):422–427, 1998.

[41] Mukund Sundararajan, Ankur Taly, and Qiqi Yan. Axiomatic attribution for deep networks. In Proceedings of the 34th International Conference on Machine Learning-Volume 70, pages 3319–3328. JMLR. org, 2017.

[42] Brandon Carter, Jonas Mueller, Siddhartha Jain, and David Gifford. What made you do this? understanding black-box decisions with sufficient input subsets. arXiv preprint arXiv:1810.03805, 2018.

[43] Brandon Carter, Maxwell Bileschi, Jamie Smith, Theo Sanderson, Drew Bryant, David Belanger, and Lucy Colwell. Critiquing protein family classification models using sufficient input subsets. In ICML Workshop on Computational Biology, 2019.

[44] Kaiming He, Xiangyu Zhang, Shaoqing Ren, and Jian Sun. Deep residual learning for image recognition. In Proceedings of the IEEE conference on computer vision and pattern recognition, pages 770–778, 2016.

[45] Fisher Yu and Vladlen Koltun. Multi-scale context aggregation by dilated convolutions. arXiv preprint arXiv:1511.07122, 2015.

[46] Sepp Hochreiter and Jürgen Schmidhuber. Long short-term memory. Neural computation, 9(8):1735–1780, 1997.

[47] Norman P P Jouppi, Cliff Young, Nishant Patil, David Patterson, Gaurav Agrawal, Raminder Bajwa, Sarah Bates, Suresh Bhatia, Nan Boden, Al Borchers, et al. In-datacenter performance analysis of a tensor processing unit. In 2017 ACM/IEEE 44th Annual International Symposium on Computer Architecture (ISCA), pages 1–12. IEEE, 2017.

[48] Erik LL Sonnhammer, Sean R Eddy, and Richard Durbin. Pfam: a comprehensive database of protein domain families based on seed alignments. Proteins: Structure, Function, and Bioinformatics, 28(3):405–420, 1997.

[49] Sean Eddy and Nick Carter. easel/esl-distance, 2017. URL https://github.com/EddyRivasLab/easel/blob/master/esl_distance.tex.

[50] Sean Eddy. Hmmer user’s guide. biological sequence analysis using profile hidden markov models. 2003.

[51] Panagiotis D Vouzis and Nikolaos V Sahinidis. Gpu-blast: using graphics processors to accelerate protein sequence alignment. Bioinformatics, 27(2):182–188, 2010.

[52] Samuel Ferraz and Nahri Moreano. Evaluating optimization strategies for hmmer acceleration on gpu. In 2013 International Conference on Parallel and Distributed Systems, pages 59–68. IEEE, 2013.

[53] Diederik P Kingma and Jimmy Ba. Adam: A method for stochastic optimization. International Conference on Learning Representations, 2014.

[54] Razvan Pascanu, Tomas Mikolov, and Yoshua Bengio. On the difficulty of training recurrent neural networks. In International Conference on Machine Learning, pages 1310–1318, 2013.

[55] Jan Chorowski, Dzmitry Bahdanau, Kyunghyun Cho, and Yoshua Bengio. End-to-end continuous speech recognition using attention-based recurrent nn: first results. arXiv preprint arXiv:1412.1602, 2014.

[56] Daniel Golovin, Benjamin Solnik, Subhodeep Moitra, Greg Kochanski, John Karro, and D Sculley. Google vizier: A service for black-box optimization. In Proceedings of the 23rd ACM SIGKDD International Conference on Knowledge Discovery and Data Mining, pages 1487–1495. ACM, 2017.

