## Supplementary Information for "Using Deep Learning to Annotate the Protein Universe"

### Pfam-seed random split dataset statistics

| | Number of examples | Number of families |
|---|---|---|
| Train | 1086741 | 17929 |
| Dev | 126171 | 13071 |
| Test | 126171 | 13071 |

Table 1: **Number of training and testing examples for the randomly split Pfam-seed data.** Note that 16755 families have sequences in the dev and test sets for the Pfam-full data.

### Performance of ensemble elements

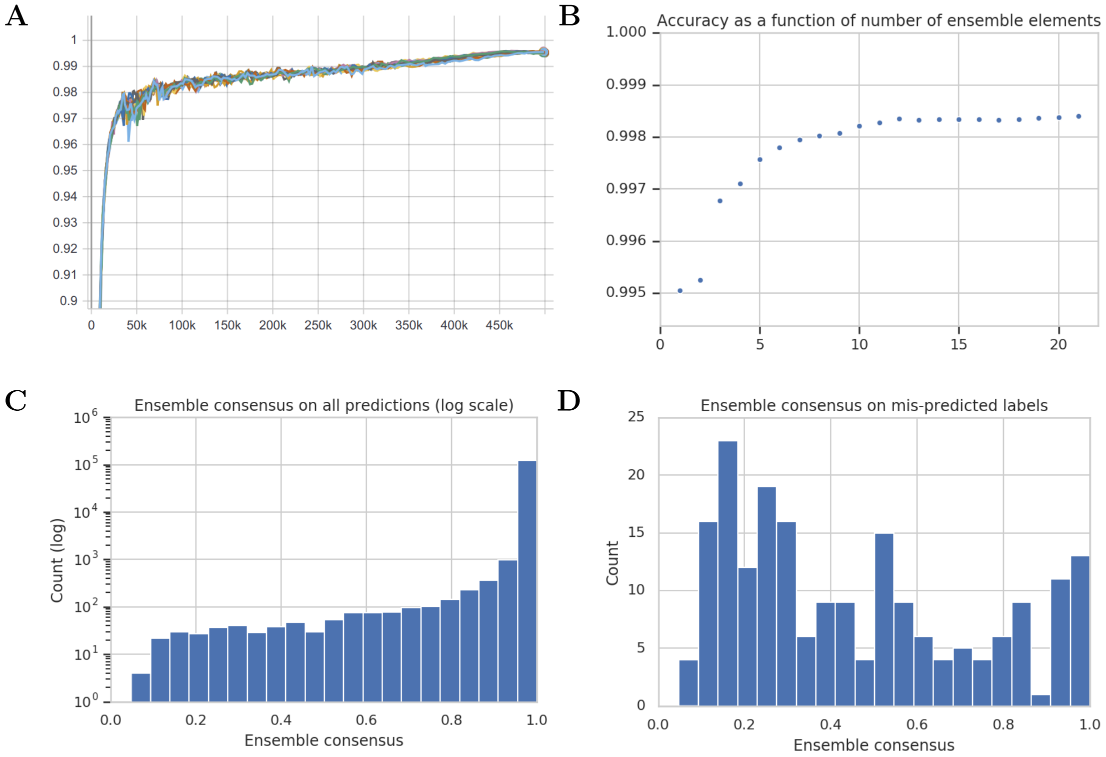


Figure 1: (A) Accuracy of training many replicates as ensemble elements on the Pfam-seed training dataset, e.g. a value at 100K on the x-axis indicates the model's accuracy on the test set after seeing 100,000 training minibatches. (B) Predictive accuracy on the held out Pfam-seed test data as a function of the number of ensemble elements. (C, D) Histogram of percentage agreement on predicted element among all ensemble elements for (C) all predictions, and (D) incorrect predictions.

Supplementary Fig. 1B shows the rapid increase in accuracy at sequence classification for the Pfam-seed dataset as a function of the number of ProtENN elements, with saturation at about 13 models. The training of our neural networks was subject to sources of stochasticity including variable initializations, example ordering, and floating point computations on GPUs. As such, a natural question to ask is "How repeatable is the training of these neural networks?" The accuracy is very stable: changing model hyperparameters did not have a large effect on the accuracy achieved. The accuracy of multiple ensemble elements (replicates) with identical hyperparameter configurations is shown in Supplementary Fig. 1B. As reported in the main text, the actual sequences that were misclassified were less stable, leading to the ProtENN ensemble.

To describe the extent to which the predictions of each model in ProtENN agree, we calculate the ensemble consensus as the ratio of votes for a particular label divided by the number of ensembled elements. We report this statistic for both misclassified sequences, and over the entire test dataset in Supplementary Fig. 1C and D. The ensemble elements generally agree with each other, though there is a nontrivial tail of disagreement, which sometimes results in an incorrect prediction.

#### Uniformly misclassified sequences

| sequence name | predicted label family name | true label family name |
|---|---|---|
| R7EGP4_9BACE/13-173* | PF04893.17 YIP1 | PF06930.12 DUF1282 |
| A0A1U8BSR7_MESAU/33-212 ‡ | PF18778.1 NAD1 | PF18782.1 NAD2 |
| A0C5A2_PARTE/441-468† | PF00036.32 EF_Hand_1 | PF13202.6 EF_Hand_5 |
| B2V696_SULSY/121-156† | PF00132.24 HEXAPEP | PF14602.6 HEXAPEP2 |
| Q7UQN3_RHOBA/655-688 † | PF00515.28 TPR | PF07719.17 TPR_2 |
| Q9VRV2_DROME/636-669 † | PF00515.28 TPR | PF07719.17 TPR_2 |
| C5FK45_ARTOC/187-218 † | PF00023.30 ANK | PF13606.6 ANK_3 |
| Q9T217_BPPHC/205-280 † | PF01751.22 Toprim | PF13662.6 Toprim_4 |
| F6ZV43_XENTR/909-973 † | PF00536.30 SAM_1 | PF07647.17 SAM_2 |
| Q01NX2_SOLUE/7-54 | PF13400.6 TAD | PF07811.12 TADe |
| H0W9E2_CAVPO/1073-1095 | PF00560.33 LRR_1 | PF05923.12 APC repeat |

Table 2: The 11 test sequences that all ensemble elements classify incorrectly, in exactly the same way. $*$ This sequence was annotated as belonging to a DUF, but has since been merged into YIP. ‡ The true label in Pfam may be incorrect, and instead be the predicted label NAD2 is a family that only includes deaminases found only in amphibians, and MESAU is a hamster, while NAD1 (ProtENN's predicted label) includes a wider array of organisms. † These sequences were split into a different family, perhaps because a single HMM could not capture all elements with this function [31].

We found that there were 11 sequences that were classified incorrectly in exactly the same way by every element of the ensemble used for ProtENN, listed in Supplementary Table 2). Our analysis of these sequences suggested that there may be some ambiguity over their correct family label in each case. For example, the sequence R7EGP4_9BACE/13-173 has a 100% identity match (found in UniProtKB) to a sequence that is classified as belonging to the YIP family. Moreover, this sequence has been independently annotated with a Gene Ontology term GO:0016021 (integral component of membrane), which matches that of the

YIP family.

In a second example, the true family for Q7UQN3_RHOBA/655-688 and Q9VRV2_DROME/636-669, PF07719, has the following description: *This Pfam entry includes outlying Tetratricopeptide-like repeats (TPR) that are not matched by PF00515.* This indicates that this family was created because there were TPR domains that didn't match the HMM for PF00515. This finding is significant because ProtENN was able to correctly "smooth" some noise in the training labels, and classify this protein as a member of PF00515. In a third example, the true family description for A0A1U8BSR7_MESAU/33-212 describes that this family is only for amphibian domains, but MESAU is a not an amphibian. Moreover, the predicted family is also a Novel AID APOBEC clade 1, but allows other species than mammals. This leads us to believe that the correct label is indeed the label chosen by ProtENN.

**Number of examples per sequence identity bucket: Pfam-seed random split**

| Sequence Identity Interval | Number of Sequences |
|---|---|
| 10-20 | 292 |
| 20-30 | 3628 |
| 30-40 | 9537 |
| 40-50 | 16798 |
| 50-60 | 22662 |
| 60-70 | 28277 |
| 70-80 | 40221 |
| 80-90 | 4429 |
| 90-100 | 256 |

Table 3: Number of sequences per sequence identity bucket for random Pfam-seed split in Figure 1A.

| Sequence Identity Interval | Number of Sequences |
|---|---|
| 12-16 | 33 |
| 16-20 | 259 |
| 20-24 | 849 |
| 24-28 | 1614 |
| 28-32 | 2499 |
| 32-36 | 3569 |
| 36-40 | 4634 |

Table 4: Number of sequences per sequence identity bucket for more remote sequences of random Pfam-seed split in Figure 1B.

### Clan level analysis

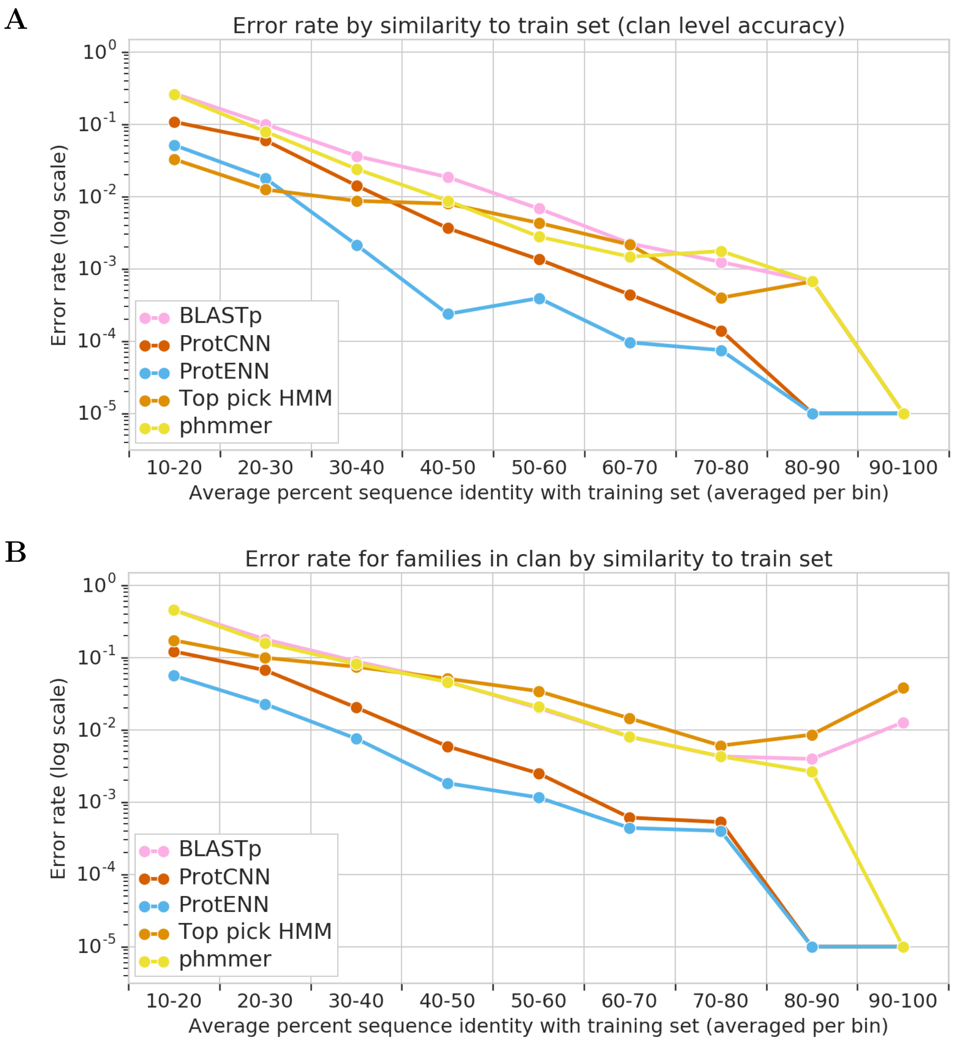


Figure 2: **Pfam Clan level analysis** (A) Clan level annotation. Held-out test error rate measured at the clan level as a function of sequence similarity to data in the Pfam-seed training set. This analysis takes account of the fact that a protein domain sequence may belong to more than one Pfam family, if the families are members of the same Pfam clan. It therefore avoids penalizing a model for annotating a sequence with a different family that belongs to the same clan. Note that the deep learning models were not retrained, and were not given any information about the existence of Pfam clans. Data has been binned into 10 bins (note some bins have more sequences than others). Under a McNemar test, the error rate of ProtENN is significantly better for sequence identity in 30-70%. All differences, both positive and negative, between ProtCNN and profile HMMs are statistically significant for sequence identity $< 70\%$. (B) Family level annotation. Here, annotation accuracy was measured at the family level for the 55604 held-out test sequences that belong to Pfam clans. Under a McNemar test at 5% significance, the reduction in error by ProtENN compared to profile HMMs is significant for sequence identity $< 90\%$.

**Multiple Family Membership**

Pfam allows a domain to belong to multiple families if these families are in the same clan [9]. Our top pick formulation of HMMer for sequence classification does not allow for multiple membership. However, within the seed sequences, there are only two sequences that belong to more than one family. The first sequence has the two distinct names ABEC3_MOUSE/245-418 and E9QMH1_MOUSE/234-407, and the second has the two distinct names NLP_DROME/6-104 and B4HZJ8_DROSE/6-104. Both of these sequences are found in our training dataset.

### Pfam-seed clustered split dataset statistics

| Sequence Identity Interval | Number of Sequences |
|---|---|
| 10-12 | 62 |
| 12-14 | 426 |
| 14-16 | 1058 |
| 16-18 | 2516 |
| 18-20 | 4419 |
| 20-22 | 6013 |
| 22-24 | 4892 |
| 24-25 | 1902 |

Table 5: Number of sequences per sequence identity bucket for clustered Pfam-seed split in Figures 2 and 4.

### Effect of family size on model performance

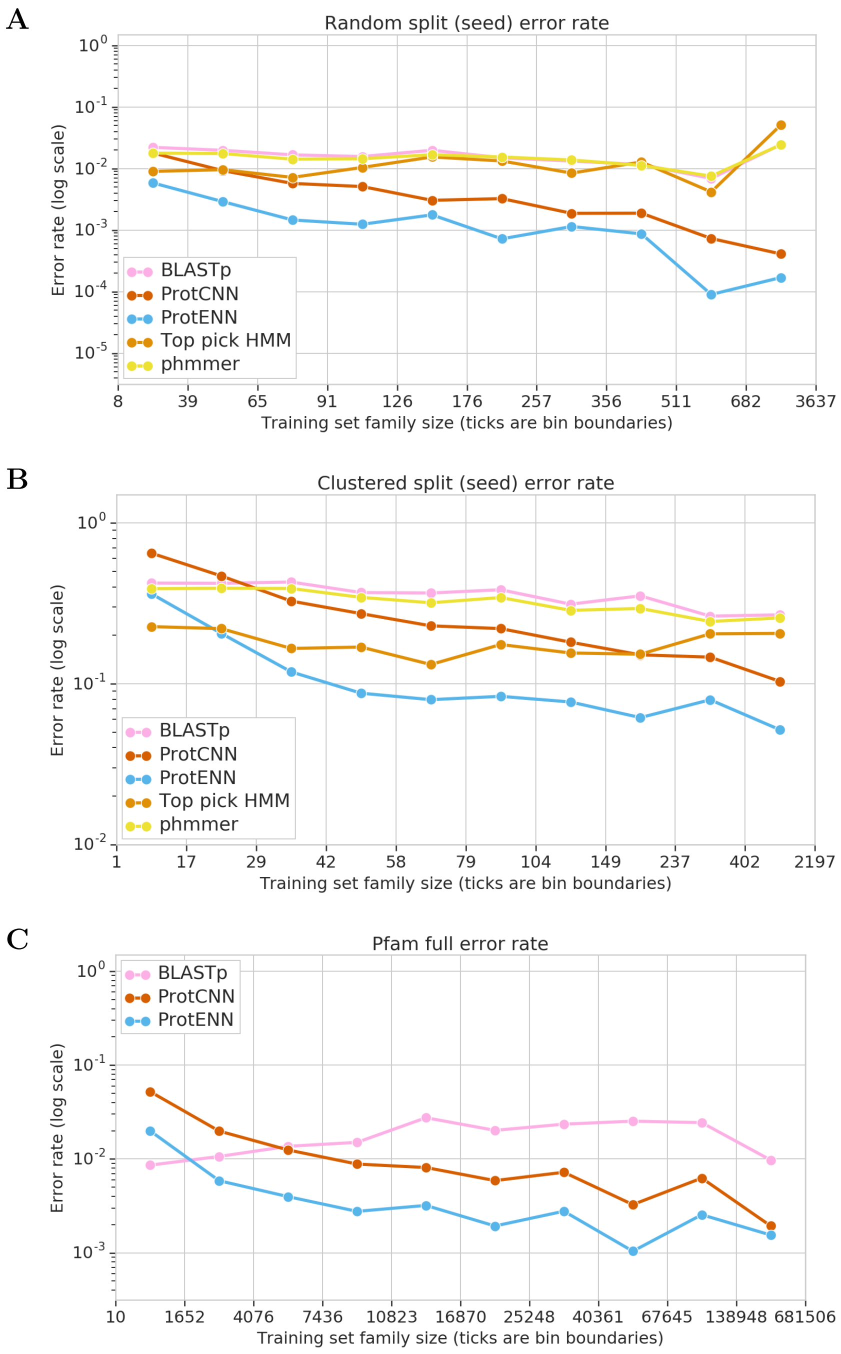


Figure 3: **Model performance stratified by family size.** (A) Pfam-seed held-out test error rate as a function of family size. Data has been binned into quantiles and the x-label ticks are the upper bound of each decile; each bin has nearly the same number of sequences. (B) Held-out test error rate as a function of training family size. X-axis ticks denote bin boundaries. Each bin has an equal number of sequences. All differences between the models, except in the bins where HMMer crosses ProtCNN and ProtENN, are statistically significant. (C) Pfam-full held-out test error rate as a function of family size, for sequences from the test set, binned in quantiles; each bin has nearly the same number of sequences.

#### Evaluation on Clustered Split using Per-Cluster Averaging

| Model | Error rate | Number of errors |
|---|---|---|
| Top Pick HMM | 15% | 3552 |
| phmmer | 31% | 6453 |
| BLASTp | 34% | 7090 |
| ProtCNN | 25% | 5117 |
| **ProtENN** | **10%** | **2174** |

Table 6: In both [29] and our work, clustering is used to split the data into train and test sets. Our construction of a test set is slightly different than that of [29], however. We first choose which clusters will be in the test set, and then include all sequences belonging to these clusters in the set. In [29], a single sequence is used for each cluster, since this helps ensure that clusters with many elements do not dominate the accuracy calculation. Here, we present results using a variant of the setup of [29]. In [29], a single sequence from each cluster is used. We instead report the expected value of this randomized procedure by first computing per-cluster average performance and then averaging these to obtain dataset-level performance. In practice, the difference between the evaluation approach in this table and in the main paper is minor because many clusters in our test set are singletons.

### Pfam-full random split dataset statistics

| Sequence Identity Interval | Number of Sequences |
|---|---|
| 20-30 | 12135 |
| 30-40 | 78075 |
| 40-50 | 182431 |
| 50-60 | 331440 |
| 60-70 | 506826 |
| 70-80 | 720693 |
| 80-90 | 1068624 |
| 90-100 | 1656757 |

Table 7: Number of sequences per sequence identity bucket for random Pfam-full split in Figure 3A.

| Sequence Identity Interval | Number of Sequences |
|---|---|
| 20-22 | 137 |
| 22-24 | 629 |
| 24-26 | 1792 |
| 26-28 | 3600 |
| 28-30 | 5977 |
| 30-32 | 8967 |
| 32-34 | 12090 |
| 34-36 | 15311 |
| 36-38 | 19530 |
| 38-40 | 22177 |

Table 8: Number of sequences per sequence identity bucket for more remote sequences of random Pfam-full split in Figure 3B.

### Few Shot Sequence Classification

As described in the main text, we construct $F$-dimensional embeddings using an *embedding network* consisting of all but the final layer of a pretrained CNN. We evaluate our embeddings in terms of their ability to provide accurate nearest-neighbor classification. When given with a new sequence, we compute its embedding and find proteins with known labels that have nearby embeddings. This approach is fundamentally different than using a CNN for classification, where we are constrained to only predict families that were seen during model training. Along the lines of popular tools such as BLAST or phmmer, we can perform nearest neighbors classification with any set of proteins and any annotation scheme, independent of how the embedding network was trained. A key difference is that we avoid direct comparisons between sequences and instead compute similarity in terms of embeddings, which uses linear algebra routines that can be accelerated substantially using modern hardware. The shared model has been trained across Pfam families, so that general protein sequence statistics need not be rediscovered from a few examples of the novel family, but instead can be used as a prior from which the statistics of the novel family can be derived.

We first train ProtCNN using the procedure described in the main text, and discard the final layer. Then, we compute the output of this embedding network for every sequence in the training set and compute a linear whitening transformation such that their covariance becomes the identity. This whitening transformation is also applied to the embeddings of test sequences, and embeddings are compared in terms of their cosine similarity.

For one shot learning, we chose the first example from the Stockholm file in each family in Pfam-seed as the representative training example for that family. For Top Pick HMM, we discarded alignment information from this sequence, and then trained and HMM using only this sequence for the small families.

For two-shot learning, we chose the first two examples. We decided not to implement two-shot Top Pick HMM so as to avoid making decisions about alignments for these two sequences.

### Computational Performance

Benchmarking for baselines was performed run on a Google Cloud Platform (GCP) n1-standard-32 instance with 32 Intel Broadwell cores, 120G RAM, and solid-state disk drive running Ubuntu Linux 16.04. `blast` and HMMER versions were 2.2.31+ and 3.2.1, respectively (note that this is the most recent release of HMMER and is different to the version benchmarked for performance). ProtCNN runtimes on CPU and GPUs were run on a n1-standard-8 (8 cores, 32 Gb, SSD) Google Cloud Platform instance with an attached NVIDIA P100 GPU:

```
gcloud beta compute --project "${PROJECT}" instances create \
"${GPU_INSTANCE}" --zone "us-west1-b" \
--machine-type "n1-standard-8" \
```

```
--image=ubuntu-1604-lts-drawfork-v20190424 \
--image-project=eip-images --boot-disk-size "250" \
--boot-disk-type "pd-ssd" --accelerator type="nvidia-tesla-p100,count=1" \
--maintenance-policy TERMINATE --min-cpu-platform "Intel Skylake"
```

In order to minimize the overhead of input and output (IO), a common bottleneck for `blast` and HMMER, all data files were stored directly in RAM in `/dev/shm` in order to eliminate, as much as possible, IO overhead. Even our largest set of sequences (54M full train) is only 9.2Gb uncompressed, less than 10% of the RAM available on the nt-standard-32 and ~30% of the RAM of the nt-standard-8 instance.

Both `hmmsearch` and `hmmscan` were allowed two cores (`--cpu` 1 argument) as recommended for its master/worker software architecture. `blast` was run with a single core (`--num_threads` 1). ProtCNN inference was run using a custom python script that (a) read in FASTA records and (b) ran inference of the ProtCNN as a TensorFlow SavedModel with command line flags to limit access to a single CPU core and/or the GPU.

All timings were run in three replicates and the user time averaged over replicates. A completely independent second machine was used by another user, producing similar runtime estimates (data not shown).

The runtime of blastp against the full train sequences database was run with 32 cores, as the blastp built-in parallelism exhibits near linear scaling with cores and, even with 32 cores end-to-end runtime is  6 hrs in our timing test.

Overall, there were 126,171 sequences in seed test sequences, which we downsampled to a 10% fraction of 12,617 sequences for runtime estimates. The seed train set has 1,086,741 sequences, full test has 5,445,307, and full train has 43,641,836 sequences.

Note that hmmsearch runs  11x faster than hmmscan for our task, so we discuss the performance of hmmsearch in the main text but the runtimes for both programs are provided for completeness.

The complete list of unix command lines needed to reproduce these times are provided below:

```
# Create the machine [On the GCP instance, run once]
PROJECT=YOUR_GCP_PROJECT
ZONE=YOUR_GCP_ZONE

gcloud beta compute --project "${PROJECT}" instances create \
"blast-hmmer-timing" --zone "${ZONE}" --machine-type "n1-standard-32"  \
--image=ubuntu-1604-lts-drawfork-v20190424 --image-project=eip-images \
--boot-disk-size "250" --boot-disk-type "pd-ssd"

gcloud beta compute ssh blast-hmmer-timing

# make a location in memory so we can run everything in the machine's memory.
```

```
TIMING_DIR=/dev/shm/timing
sudo mkdir ${TIMING_DIR}
sudo chown ${USER} ${TIMING_DIR}

# install required software
sudo apt-get --yes install make gcc

# Install blast
sudo apt-get --yes install ncbi-blast+

# install hmmer version 3.2.1
cd ~
wget http://eddylab.org/software/hmmer/hmmer-3.2.1.tar.gz
tar zxf hmmer-3.2.1.tar.gz
pushd hmmer-3.2.1
./configure --enable-threads
make
make check
popd
HMMSEARCH=~/hmmer-3.2.1/src/hmmsearch
HMMSCAN=~/hmmer-3.2.1/src/hmmscan
HMMPRESS=~/hmmer-3.2.1/src/hmmpress

# Get the PFam 32.0 hmm profiles.
cd ${TIMING_DIR}
wget ftp://ftp.ebi.ac.uk/pub/databases/Pfam/releases/Pfam32.0/Pfam-A.hmm.gz
gunzip Pfam-A.hmm

# Create the compressed hmm db for hmmscan
${HMMPRESS} Pfam-A.hmm

PROTEINS_BUCKET=gs://brain-genomics-public/research/proteins/timing

# Grab seed_train.fasta, full_train.fasta and seed_test.fasta
for f in full_train.fasta seed_train.fasta seed_test.fasta; do
  echo "Downloading file $f"
  curl -o ${TIMING_DIR}/${f} \
    https://storage.googleapis.com/brain-genomics-public/research/proteins/timing/${f};
done
```

```
wc -l *.fasta
# Expect to see 252342 lines for seed_test.fasta,
# 2173482 for seed_train.fasta
# and 87283672 for full_train.fasta.

# Create a 10% subset of the seed_test.fasta
head -n 25234 seed_test.fasta > seed_test.10_percent.fasta

# Create blast databases for seed and full train
makeblastdb -in seed_train.fasta -dbtype prot
makeblastdb -in full_train.fasta -dbtype prot


# Use the 10% sample of seed_test so the programs finish in a shorter timespan.
timing_fasta=seed_test.10_percent.fasta

# Use the full seed_test.fasta for a more complete runtime estimate.
# timing_fasta=seed_test.fasta

# We are using three replicates.
N_REPLICATES=3
HMMER_NCORES=1

# Time hmmscan and hmmsearch of seed_test.fasta against Pfam-A.hmm.
for replicate in $(seq $N_REPLICATES); do
for binary in ${HMMSCAN} ${HMMSEARCH}; do
  echo "Profiling hmmer ${binary} [replicate ${replicate}]"
  name="hmmer.${timing_fasta}.${binary##*/}.cores_${HMMER_NCORES}.rep_${replicate}"
  (time ${binary} \
    --cpu ${HMMER_NCORES} \
    --tblout ${name}.txt \
    -o ${name}.log \
    Pfam-A.hmm ${timing_fasta}) &> ${name}.time.log
  cat ${name}.time.log
done
done

# We want to use a different number of cores for each blast calculation.
# For seed, we want to use a single core so it's more directly comparable
# to hmmer. But blastp running on the 10% subset against the full
# training database takes a really long time. So we'll use all cores for that.
```

```
declare -A blast_database_ncores
blast_database_ncores[seed_train.fasta]=1
blast_database_ncores[full_train.fasta]=32

# Time blast against seed_train
for replicate in $(seq $N_REPLICATES); do
for blast_database in seed_train.fasta full_train.fasta; do
  ncores=${blast_database_ncores[${blast_database}]}
  echo "Profiling blastp against database \
  ${blast_database} with ${ncores} cores [replicate ${replicate}]"
  name="blast.${timing_fasta}.${blast_database}.cores_${ncores}.rep_${replicate}"
  (time blastp \
    -query ${timing_fasta} \
    -db ${blast_database} \
    -outfmt 10 -max_hsps 1 -num_alignments 1 \
    -num_threads ${ncores} \
    -out ${name}.out ) &> ${name}.time.log
  cat ${name}.time.log
done
done

# grep out all of the results:
fgrep real *.time.log
```

### Saturation mutagenesis experiments

| Sequence Name | Residues | Sequence |
|---|---|---|
| AT1A1_PIG | 161-352 | NMVPQQALVIRNGEKMSINAEEVVVG<br>DLVEVKGGDRIPADLRIISANGCKVD<br>NSSLTGESEPQTRSPDFTNENPLETR<br>NIAFFSTNCVEGTARGIVVYTGDRTV<br>MGRIATLASGLEGGQTPIAAEIEHFI<br>HIITGVAVFLGVSFFILSLILEYTWL<br>EAVIFLIGIIVANVPEGLLATVTVCL<br>TLTAKRMARK |
| V2R_HUMAN | 54-325 | SNGLVLAALARRGRRGHWAPIHVFIG<br>HLCLADLAVALFQVLPQLAWKATDRF<br>RGPDALCRAVKYLQMVGMYASSYMIL<br>AMTLDRHRAICRPMLAYRHGSGAHWN<br>RPVLVAWAFSLLLSLPQLFIFAQRNV<br>EGGSGVTDCWACFAEPWGRRTYVTWI<br>ALMVFVAPTLGIAACQVLIFREIHAS<br>LVPGPSERPGGRRRGRRTGSPGEGAH<br>VSAAVAKTVRMTLVIVVVYVLCWAPF<br>FLVQLWAAWDPEAPLEGAPFVLLMLL<br>ASLNSCTNPWIY |

Table 9: Wildtype sequences, keyed by Uniprot ID, that were used for saturation mutagenesis predictions.

As reported in the main text, we challenged ProtENN trained on the Pfam-full dataset to distinguish between single amino acid variants of protein domain sequences. Using the above method, we examined the purported helical propensity of each amino acid in the transmembrane regions (the amino acids whose substitutions least changed the predicted function of the domain), and found that the list, from most to least favored, is `VLIAM FTWCY SNGQH PRDKE`. The fact that the charged amino acids, along with proline, are the least favored is in accord with our understanding of 1) the unfavorability of polarity in the transmembrane region; 2) the sharp binding angle of proline. We also note the visual effect of substituting glycine is much like that of substituting proline: we know that glycine has low helical propensity [40]. For this figure and for that of the main text, we clip large values to show fine-grained color differentials. The wild-type sequences used for Figures 5B and Supplementary Fig. 4 are shown in Table 9.

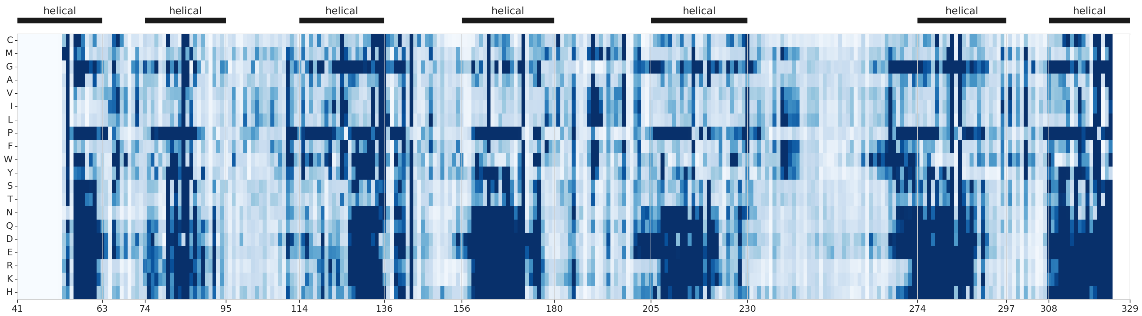


Figure 4: Predicted change in function for each missense mutation in vasopressin domain V2R_HUMAN/54-325 from family PF00001.21. The x-axis is residue indices in the protein P30518 (the domain starts at index 54), the y axis is the substitution of a particular amino acid, and a dark color saturation describes a large predicted change in function.
The model (trained on Pfam-full) appropriately predicts that substituting proline, glycine, or charged amino acids in the transmembrane helix regions is very likely to change the function of the protein substantially.

| | Number of examples | Number of families |
|---|---|---|
| Train | 1296280 | 17929 |
| Dev | 21510 | 4323 |
| Test | 21293 | 3097 |

Table 10: **Number of training and testing examples for the clustered split of the Pfam-seed data.**

### Benchmark Dataset

Pfam contains one profile HMM for each family, constructed from the family seed alignment. These manually curated seed alignments contain between 1 and 4545 sequences, originally picked due to their trusted functional annotations [48], avoiding some of the circularity of using sequence labels assigned by a profile HMM model. However, many Pfam seeds have recently been rebuilt to incorporate sequences from Pfam-full that were found by the original family HMM, reducing their model-free status [33]. Pfam-seed sequences vary between 4 and 2037 amino acids in length, with 27045 seed sequences of length $> 500$. Supplementary Fig. 5 contains a histogram of Pfam-seed family sizes, the Pfam-seed sequence length distribution and also the frequency of amino acid usage in the Pfam-seed dataset.

Family-specific gathering thresholds, shown in Supplementary Fig. 5D, are used by HMMer 3.1b to determine whether a sequence belongs to each family [9]. The role of these gathering threshold is to increase coverage and decrease false positives. However our setup simply takes the top match by score, regardless of the assigned gathering threshold. This

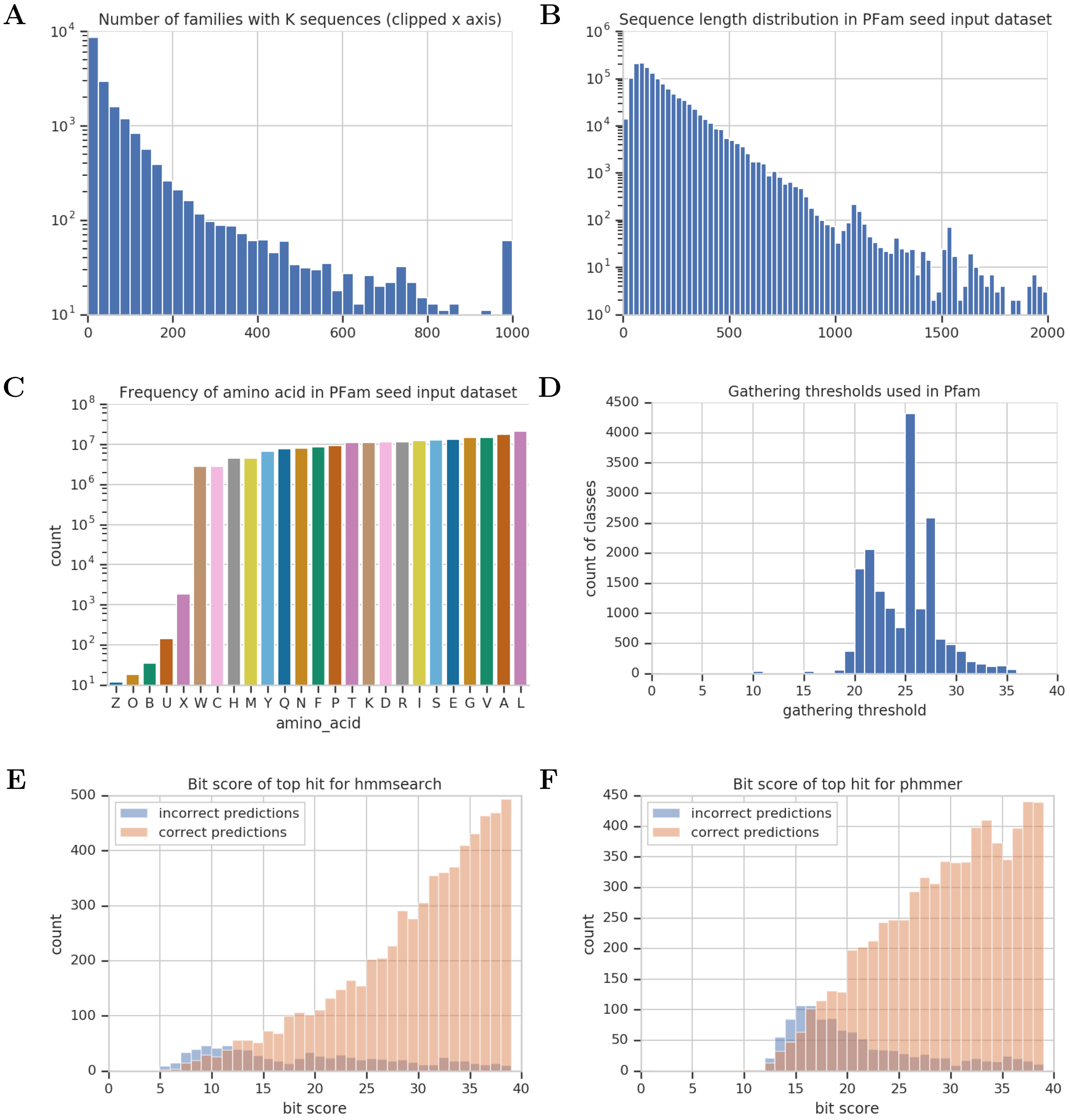


Figure 5: Benchmark Pfam-seed dataset statistics calculated across the entire Pfam-seed dataset. (A) Number of sequences per family. Values larger than 1000 are clipped to the last histogram bucket. (B) Sequence length distribution of unaligned sequences. (C) Frequency of amino acids in sequences in the dataset. (D) Histogram of gathering thresholds used in Pfam 32.0. Scores achieved by the top hits for (E) the HMM top pick model and (F) phmmer; the x axis has been truncated in both panels. Compare to (D) the actual histogram of gathering thresholds used by Pfam.

raises the question of whether implementing the family-specific gathering thresholds would have improved the accuracy score achieved by the HMM top pick algorithm.

To address this, Supplementary Fig. 5E and F shows the distribution of top scores for both HMMer and phmmer, for both correct and incorrect predictions. We note that the majority of incorrect predictions have scores *below* the assigned gathering thresholds. However, there also appear to be at least as many correct predictions below these values. This qualitative analysis is backed up by experimentation wherein we determined 8.5% of top picks were below their family-specific gathering thresholds from Pfam. As such, using the assigned gathering thresholds would not have helped performance, and effective use of gathering thresholds would have required us to re-tune each of these values for the training dataset used in this benchmark study.

### Details of Neural Network Architectures and Training

In a residual network (ResNet) [44], the layers are built up additively, with $f_i = f_{i-1} + g_i(f_{i-1})$. Here, each $f_i$ is an $L \times F$ array and $g_i(\cdot)$ is an additional one-layer convolutional network (along with a kernel-size-one bottleneck convolution; see Figure 6C) with trained weights specific to that layer. In our model, $f_0$ is obtained by a convolutional layer with $F$ channels applied to the output of the input network, with no bottleneck convolution applied before the residual blocks. Each ResNet layer maintains a $L \times F$ representation; no downsampling is performed until the final pooling step. We also note that a convolutional layer is used before any residual block, so as to convert the per-residue representation into the correct shape before consumption by the residual blocks.

We primarily focus on convolutional neural networks (CNNs) to construct this $L \times F$ array, since they are fast to train and evaluate on modern hardware, an advantage that is even more pronounced when evaluating large sets of sequences in parallel. Convolutional architectures are also easily composed into higher-order interactions. The $L \times F$ array is then pooled along the length of the sequence, ensuring invariance to padding. Hyperparameters tuned for each neural network include the choice of $F$ and max vs mean pooling in addition to network depth, which was varied between 1 and 6 layers.

### Dilated Convolutions

Dilated convolutions are a popular method for enabling CNNs to capture long range interactions across the inputs [45]. One way to to model these long-range interactions would be to use convolutions with very wide kernels. However, doing so increases the computational complexity of prediction and introduces a considerable number of parameters to train. Instead, dilated convolutions use convolution kernels with holes in them, so that the complexity and number of parameters is the same as small, local convolutions, but the overall receptive field of the convolution is wide.

Consider a convolution with kernel width 5, and let $f_{i,j}$ be the representation in layer $i$ of the CNN at position $j$ in the sequence. In a traditional 1-dimensional convolution, $f_i$ is a linear function of

$$\{f_{(j-2)}, f_{(i-1),(j-1)}, f_{(i-1),j}, f_{(i-1),(j+1)}, f_{(i-1),(j+2)}\}.$$

In a dilated convolution with dilation rate $r$, it is a function of

$$\{f_{(i-1),(j-2r)}, f_{(i-1),(j-r)}, f_{(i-1),j}, f_{(i-1),(j+r)}, f_{(i-1),(j+2r)}\}.$$

At each layer of our CNN, $r$ is increased by a factor of $k$, so the overall receptive field size of the CNN is exponential in its depth. Specifically, if the model has $n_1$ non-dilated layers followed by $n_2$ dilated layers, $k$ is the kernel width and $r$ the dilation rate, then the receptive field size is $k + 2(k-1)(n_1 - 1) + 2(k-1)\sum_{i=1}^{n_2} r^i$. These terms correspond to the first layer, the remaining non-dilated layers, and the dilated layers respectively. Supplementary

Fig. 1 illustrates the relationship between receptive field size and classification accuracy of the resulting ProtCNN model for the Pfam-seed dataset.

#### Model Invariance to Padding

At both train and test time, our model processes sequences in batches. The batches are of variable length, so input one-hot sequences are padded with zeros before being stacked together in a rectangle that can be processed in parallel on a GPU (see Fig. 6C). It is imperative that our model's predictions are insensitive to the padding, as the amount of padding depends on the other sequences in the batch (we pad to the longest sequence in the batch). For RNNs, this can be achieved by ending the recurrence of the RNN at the end of the un-padded sequence. For CNNs, our model maintains an $L \times F$ array of features at every layer, where each column corresponds to a specific location in the input sequence. Before each convolution or batch normalization operation, we zero-out the features in any location that corresponds to padding in the input sequence. This ensures that the model's predictions are insensitive to padding at test time. However, the dynamics of training our CNNs are still effected by padding, since batch normalization computes feature averages across the length of the sequence, and these lengths vary due to padding.

### Model Training

We use the Adam optimizer [53]. The learning rate is subject to exponential decay following a warm-up period, and the length of the period was not treated as a tunable hyperparameter. At train time, we present the model with randomly-drawn batches. Consistent with popular experience [54], we find that it is important to use gradient clipping for the RNN, and adaptive gradient clipping worked significantly better than static gradient clipping [55], so we use adaptive gradient clipping for all the deep models.

### Neural Network Hyperparameters

| Model Type | Hyperparameter | Search Range |
|---|---|---|
| ProtCNN | batch size | 32, 64, 128, 256 |
| | dilation rate | 1, 2, 3, 5 |
| | filters | 300 thru 3000, increments of 100 |
| | ResNet block of first dilated layer | 2, 3 |
| | kernel size | 3, 7, 9, 11, 21, 31 |
| | ResNet layers | 1 thru 6 |
| | learning rate | 1e-05, 5e-05, 1e-4, 5e-4, 1e-3 |
| | learning rate decay steps | 1e3,1e4,1e6, decay off |
| | pooling | max, mean |
| 1, 2 ResNet block CNN | dilation rate | 1, 2, 3, 5 |
| | filters | 150 thru 500, increments of 50 |
| | ResNet block of first dilated layer | 2, 3 |
| | kernel size | 3, 7, 9, 11, 21, 31 |
| | learning rate | 1e-4, 5e-4, 1e-3 |
| | pooling | max, mean |
| RNN | learning rate | 1e-4, 5e-4, 1e-3 |
| | number of hidden units | 25 thru 2048, increments of 1 |
| | pooling | max, mean |
| kmer | embedding rank | 100, 1000, 10000 |
| | learning rate | 1e-4, 5e-4, 1e-3 |
| | ngram order | 1 thru 5 |

Table 11: Search ranges for hyperparameter values, by model.

Our embedding network architectures involve a variety of hyperparameters as outlined in Supplementary Tables 11 and 12. For all networks, the "dev" fold is used to identify the optimal hyperparameter settings, while model performance statistics are reported using the completely distinct "test" fold. The CNN hyperparameters are tuned using values sampled at random from each hyperparameter search range, reported in Supplementary Table 11. The number of searched values is reported in Supplementary Table 13. We carried out an initial study that identified the most promising architecture. Supplementary Fig. 1B illustrates hyperparameter tuning.

We studied the impact of different hyperparameter settings on ProtCNN. As shown in Supplementary Table 11 we also allowed the batch size to vary, and introduced additional

| | ProtCNN | 2 Block CNN | 1 Block CNN | RNN |
|---|---|---|---|---|
| batch size | 32 | 64 | 64 | 64 |
| dilation rate | 3* | 2* | | |
| filters | 1100* | 500* | 500* | |
| first dilated layer | 2* | 2* | | |
| gradient clip | 1 | 1 | 1 | 1 |
| kernel size | 9* | 31* | 31* | |
| Number hidden units | | | | 1244* |
| learning rate | .0001* | .001* | .0001* | .0005* |
| learning rate decay rate | 0.997 | 0.997 | 0.997 | 0.997 |
| learning rate decay steps | 1000* | 1000 | 1000 | 1000 |
| learning rate warmup steps | 3000 | 3000 | 3000 | 3000 |
| number of ResNet layers | 5* | 2 | 1 | |
| pooling | max* | max* | max* | mean* |
| ResNet bottleneck factor | 0.5 | 0.5 | 0.5 | |
| train steps | 500000** | 400000 | 400000 | 300000 |

Table 12: Hyperparameters used in neural networks for Pfam-seed dataset. Column headers are model-type. An asterisk denotes a tuned value. Two asterisks denote that the model was overfit, and the number of tuning steps was chosen post-hoc so as to maximize dev-set performance.

| Model Type | Search Algorithm | Approx. number of samples |
|---|---|---|
| CNN (all depths) | random sampling | 17000 † |
| RNN | Gaussian process | 250 |
| kmer | random sampling | 50 |

Table 13: Search algorithms and number of samples for hyperparameter tuning, by model. † Many of these configurations were not feasible, as they did not fit in GPU memory.

learning rate decay parameters in this study. Moreover, the number of filters was greatly increased, and the number of layers was allowed to vary as a hyperparameter. These modifications helped to maximize the performance of ProtCNN in terms of accuracy. However, they made the resulting model more difficult to interpret, in the sense that it became difficult to attribute increases in performance to specific parameters such as the size of the receptive field.

| | ProtCNN |
|---|---|
| batch size | 64 |
| filters | 2000* |
| first dilated layer | NA |
| gradient clip | 1 |
| kernel size | 21* |
| learning rate | .001* |
| learning rate decay rate | 0.997 |
| learning rate decay steps | 1000* |
| learning rate warmup steps | 3000 |
| pooling | max* |
| ResNet bottleneck factor | 0.5 |
| train steps | 1100000** |

Table 14: Hyperparameters used in neural networks for Pfam-full dataset. An asterisk denotes a tuned value. Two asterisks denote that the model didn't necessarily converge, but was ended after a reasonable time training (17 days).

Supplementary Table 14 shows the ProtCNN hyperparameters used for the Pfam-full dataset. For the RNN models, we tuned the hyperparameters jointly using Bayesian optimization with a Gaussian process regressor [56]; see Supplementary Table 11 for the search ranges, Supplementary Table 12 for the values, and Supplementary Table 13 for the number of searched values. RNNs using the final sequence-dimension LSTM cell output could only predict the mode of the distribution, however using the mean or max across the sequence dimension fixed this inability to propagate features through time. Increasing the number of layers did not improve performance, additional RNN hyperparameter settings are provided in Supplementary Table 12. It was hard to find RNN configurations with stable training dynamics, and RNNs took substantially longer to train. Furthermore, while CNN computations can be parallelized along the length of the sequence, RNN computations must be done sequentially, resulting in considerable slowdown for long sequences.

| | kmer |
|---|---|
| batch size | 64 |
| gradient clip | 1 |
| learning rate | .0005* |
| learning rate decay rate | 0.997 |
| learning rate decay step | 1000 |
| learning rate warmup steps | 3000 |
| kmer order | 2* |
| number of hash buckets | 10000* |
| train steps | 300000 |

Table 15: Hyperparameters used in kmer benchmark models. An asterisk denotes a tuned value.

### Comparison of Sequence Identity Calculations

For Pfam-seed we use the Pfam family alignment to compute a pairwise distance between every held-out test sequence and sequences from the same family that are contained in the training set. Another metric of distance between each held-out test sequence and the training set is provided by the percent sequence identity measured by BLASTp. To provide an idea of the differences between these metrics, Supplementary Fig. 6 compares them across all 126171 sequences contained in the randomly split Pfam-seed held-out test set.

For the split of Pfam-full, we observe an increase in model error rate for BLASTp in the last decile of pairwise sequence identity computed using BLASTp (see Fig. 3A). There are two potential sources for this reduction in accuracy. The first is sequences that are closer in terms of sequence identity to a member of a different family than to their own. The second is that some sequences in the dataset are sub-sequences of others. Where the sub-sequence is in the test set, BLASTp measures "100%" sequence identity with the super-sequence contained in the training set. Discerning the correct classification in these cases can be quite difficult. For example, in Pfam-full, one of the test sequences is A0A010NMM2_9MICC/241-409, and one of the training sequences is A0A010NMM2_9MICC/4-495. In this case, the former sequence has is identical to part of the latter, but it is classified differently by Pfam: the test sequence is the NAD binding domain of AdoHcyase, while the latter is the full AdoHcyase domain. This may explain some of the difficulty that BLASTp has with sequences that are very similar to those in the training set.

A

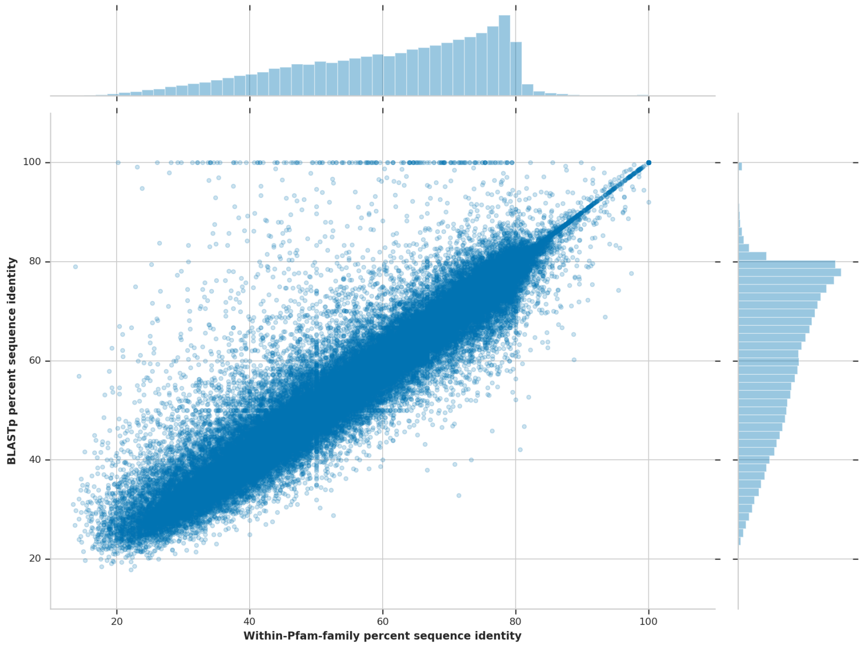


B

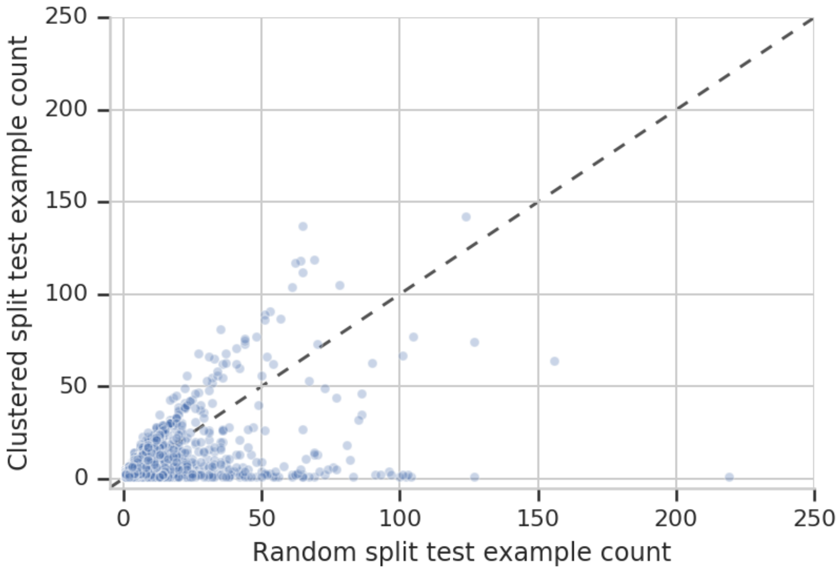


Figure 6: (A) Comparison of the within Pfam family distance calculated using the formula given in the text with the BLASTp percent sequence identity for each of the 126171 held-out test sequences of the Pfam-seed dataset. (B) Number of test-set sequences for each family in the random and clustered splits. While the clustering process is desirable because it ensures separation between train and test data, it introduces a distribution over families in the test data that is significantly different than the overall distribution in Pfam.
